# Actin-related ACTL8 localizes to the nucleus and modulates locus-specific transcription through chromatin regulation

**DOI:** 10.64898/2026.08.21.746286

**Authors:** Nuria Cortes-Silva, Sanjana Banerjee, Kristen Dominique Amarillo, Sahiti Peddibhotla, Isabel Mejia Natividad, Betta Chopard, Christopher B. Geyer, Janet M. Young, Sandipan Brahma, Courtney M. Schroeder

## Abstract

Actin-related proteins (Arps) are evolutionarily ancient and broadly expressed, yet some Arps evolved in mammals and acquired germline-specific expression. Here, we investigate the germline-specific Arp “Actin-like 8” (ACTL8), which is also misexpressed in multiple cancers with unexplored molecular roles. We show that ACTL8 surprisingly localizes to the nucleus of spermatogonia in human testes and breast and lung cancer cells. In breast cancer cells, ACTL8 misexpression alters transcriptional programs primarily in cell migration pathways, and ACTL8 depletion increases migration rate. Mechanistically, we find that ACTL8 interacts with KDM5B, an H3K4 demethylase and transcriptional repressor, and demonstrate that this interaction contributes to changes in migration. Consistent with this, ACTL8 loss increases H3K4me3 at genes associated with migration, indicating transcriptional activation. These findings establish ACTL8 as a novel nuclear Arp that plays a role in KDM5B-mediated transcriptional repression, providing insight into its function in chromatin regulation in cancer and potentially spermatogenesis.

## Introduction

Actin is ancient and found across the tree of life with essential roles, including in cell motility and cell division^1,2^. In the early 1990s, it was discovered that eukaryotes encode not only for actin but also actin-related proteins (Arps), collectively forming the Arp superfamily. The evolutionary history of Arps suggests that these proteins duplicated before the last common ancestor of eukaryotes and established approximately 10 subfamilies (with two appearing only in fungal species)^1,2^. Arps are considered “actin-related” primarily due to the retention of the canonical actin tertiary structure^1–3^. However, they have diverged in sequence from canonical actin, sharing ∼30% to 70% sequence identity with actin^1,2^. Their sequence divergence has resulted in distinct cellular functions, many of which are essential. Arps are expressed across most cell types and carry out roles ranging from microtubule-based transport (Arp1 and Arp10)^4–6^, actin polymerization (Arp2 and Arp3)^7^, chromatin remodeling (Arp4, Arp5, Arp6 and Arp8)^8–14^, and histone acetylation (Arp4)^10,15^. Each Arp subfamily is highly conserved and has been retained across eukaryotic evolution for their critical cellular roles.

Arps play roles in the cytoplasm or the nucleus, and a subset play roles in both contexts. The Arps that compose chromatin remodeling complexes are primarily nuclear, while Arp1 and Arp10 appear to be strictly cytoplasmic, as they are components of the dynactin complex that regulates the motility of the microtubule motor dynein^4–6^. Arp2/3 and actin, while enriched in the cytoplasm, also have important nuclear functions. Actin is found as a monomer in complex with nuclear Arps within chromatin remodeling complexes and plays roles in transcription, replication, and DNA repair together with Arp2/3^16,17^. For example, DNA double-stranded breaks trigger Arp2/3-catalyzed actin polymerization, which facilitates the movement of breaks to the nuclear periphery for repair ^18,19^. Furthermore, most Arps carry out their roles as monomers within a complex (e.g. Arps4-8), though Arp1 can polymerize like actin^20^. In sum, the Arp superfamily plays a wide range of subcellular roles.

In addition to canonical Arps, a subset of Arps has recently evolved in mammals by way of gene duplication ^21–24^. Unlike actin and canonical Arps, these non-canonical Arps frequently display tissue-restricted expression, particularly in the germline^23–27^. Mammals encode at least eight non-canonical Arps whose expression is enriched in the testis^21–24,28^. Several of these Arps localize to a unique testis-specific actin structure in post-meiotic spermatids known as the acroplaxome that regulates spermatid head reshaping and tethers the nucleus to the acrosome, which is critical for sperm-egg fusion^24,29,30^. Interestingly, the emergence of non-canonical Arps extends beyond mammals, as *Drosophila* species also encode germline-enriched non-canonical Arps^31–33^. These Arps have diversified from their canonical counterparts in both sequence and function, similar to the divergence that occurred during the ancient origin of the Arp superfamily. One non-canonical Arp in *D. melanogaster* acquired a critical role in fertility and embryonic development^32^. Together, these observations reveal that non-canonical Arps emerged independently multiple times during evolution across lineages and frequently acquired specialized germline roles. While the biological importance of some of these Arps is clear, the molecular roles of their biology are poorly understood.

Actin-like 8 (ACTL8) is a non-canonical Arp normally expressed in the testis^34,35^, though its function remains unclear. Most studies of ACTL8 describe its misexpression in cancer as a cancer/testis antigen (CTA)^36–38^. CTAs are a class of proteins normally expressed in the germline but become aberrantly expressed in cancer^39,40^. ACTL8 is misexpressed in numerous cancer types, including triple negative breast cancer^41,42^, colorectal cancer^43^, oral squamous cell carcinoma^44^, lung cancer ^45,46^, head and neck squamous cell carcinoma^28^, and gastric cancer^47^. Increased *ACTL8* expression often correlates with poor patient prognosis^28,41,43,44^. Studies across these cancer types have indicated that ACTL8 expression promotes cell proliferation, migration and invasion, which is partially due to effects on cell cycle signaling pathways in some cancers^41,42,47^. Despite its potential impact on cancer cell biology and its unique restricted expression in the testis, the molecular roles and even cellular localization of ACTL8 remain unclear in cancer and the germline.

Here, we gain insight into human ACTL8 function by combining evolutionary analyses, cytology, and gain- and loss-of-function experiments in breast cancer cells. We find that ACTL8 surprisingly localizes to the nucleus of both early male germ cells and cancer cells, identifying ACTL8 as a novel nuclear Arp. Transcriptome analyses of breast cancer cells reveal that *ACTL8* alters the expression of genes involved in cell proliferation and migration, and in support of these findings, loss of *ACTL8* accelerates cell migration. We further discovered that ACTL8 interacts with KDM5B, a histone H3 lysine 4 demethylase that functions as a transcriptional repressor^48^. To determine whether the ACTL8-KDM5B interaction indeed alters migration, we identify the minimal ACTL8-binding region of KDM5B and use it as a competitive inhibitor; indeed, overexpression of this fragment increased cell migration in a similar manner to loss of *ACTL8*. We further map H3K4me3 levels in breast cancer cells and find that upon reduced *ACTL8* expression, H3K4me3 increases at genes implicated in cell migration, indicating that ACTL8 loss enhances loci-specific transcriptional activation. Together, our findings provide key molecular insights into the function of ACTL8 and establish it as a nuclear Arp participating in KDM5B-mediated transcriptional repression. While our findings demonstrate how ACTL8 impacts cancer cell biology, they also suggest that ACTL8 evolved a specialized nuclear function in shaping the chromatin landscape in early sperm development.

## Results

### ACTL8 is a germline-specific Arp that evolved in mammals

*ACTL8* is encoded in humans, but it is unknown whether it predates primates and how conserved it is across species. We surveyed the syntenic location of *ACTL8* in well-assembled genomes of representative vertebrate species, and we identified *ACTL8* only in mammals (**Fig. 1A, Supplementary Fig. 1A, Supplementary Data 1**). Most placental mammals encode *ACTL8* except for some rodents, including mouse and rat (**Fig. 1A**). The marsupial opossum and monotreme species platypus also encode syntenic *ACTL8* orthologs, albeit notably diverged with both having ∼33% protein identity with human ACTL8. Species that did not appear to have *ACTL8* in the syntenic locus were further surveyed using tBLASTn^49^. We did not uncover *ACTL8*-like genes in non-mammalian genomes, with top tBLASTn hits being canonical Arps (**Fig. 1A, Supplementary Data 1**). Overall, we conclude that ACTL8 evolved at least 180 million years ago^50^ and appears to be exclusively a mammalian Arp.

**Figure 1:**
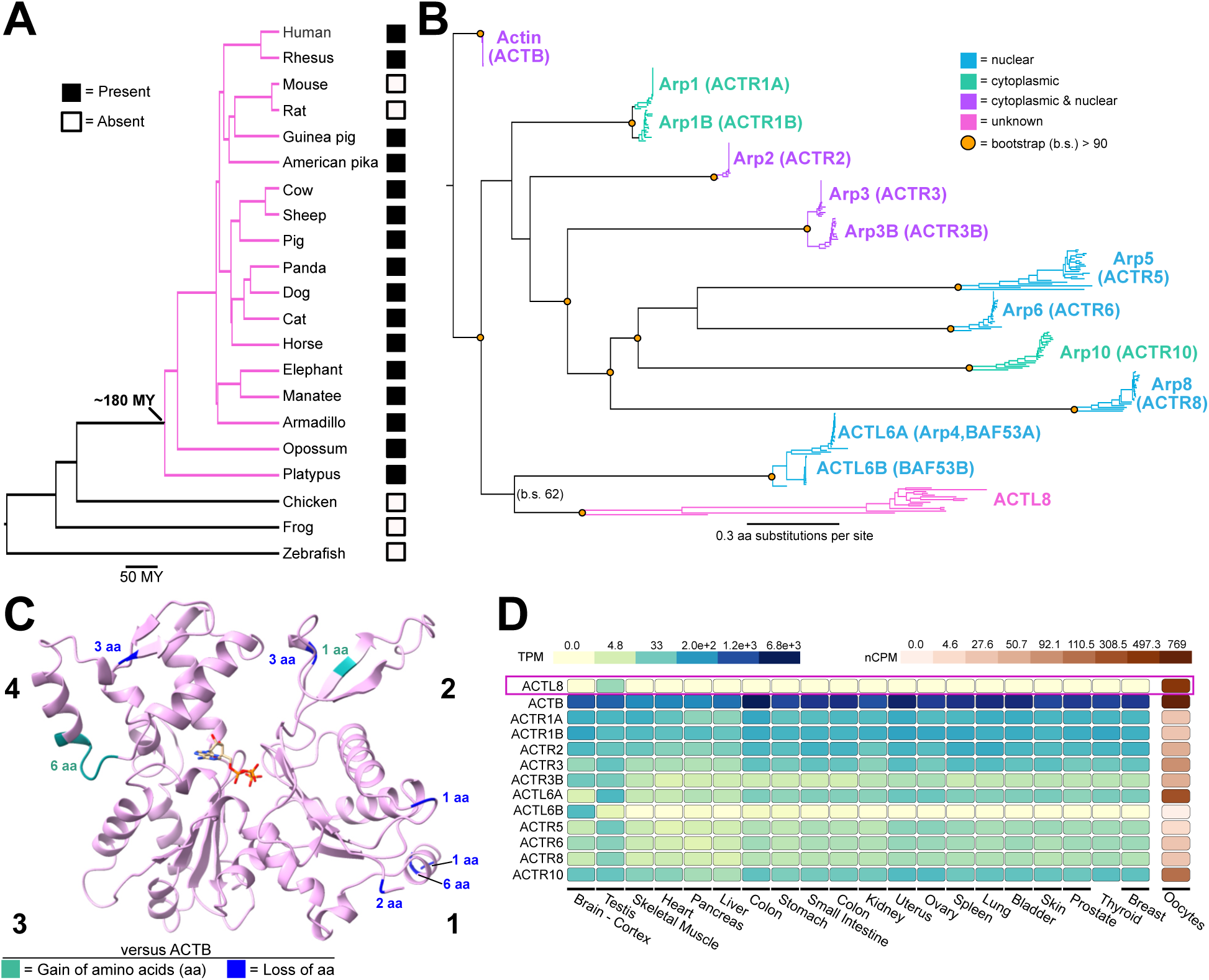
ACTL8 is a germline-specific Arp that evolved in mammals. **A)** Species tree showing presence and absence of syntenic ACTL8 orthologs. Branches including placental mammals, the marsupial opossum, and the monotreme platypus are highlighted in pink with the approximate origin of ACTL8 noted (180 MYA). **B)** Protein sequence tree including canonical Arps and ACTL8 sequences from a subset of vertebrate species. Subfamilies are color coded based on subcellular localization, and only nodes with bootstrap support greater than 90% are noted with orange circles (bootstrap support noted for node shared between ACTL8 and ACTL6A/B subfamilies). Scale is 0.3 aa substitution per site. **C)** AlphaFold 3.0^51^ predicted structure of ACTL8 bound to ATP. Subdomains of actin fold are labeled 1-4. Residues that were gained with respect to human actin (ACTB) are in green, and one residue at site of residue loss (relative to ACTB) is noted with number of lost residues. **D)** Normalized RNA-seq data for all canonical Arps and ACTL8 across healthy adult human tissues, with dark blue being the highest expression, from the Adult Genotype Tissue Expression (GTEx) Project^131^ (data obtained 5/10/26). Oocyte expression was obtained from The Human Protein Atlas^54,55^ and shown with a separate scale (dark red being highest).

We investigated the evolutionary relationship of orthologous ACTL8 protein sequences to canonical Arps across vertebrate species by conducting maximum likelihood phylogenetics analyses. We included canonical Arps from the same species that encode ACTL8 and vertebrates that do not encode ACTL8 (e.g., chicken, *Xenopus tropicalis* and zebrafish). We found that within the Arp superfamily, ACTL8 forms its own subfamily with high bootstrap support, including the diverged opossum and platypus ACTL8 orthologs (100% bootstrap, **Fig. 1B**). While the ACTL8 subfamily closely groups with the nuclear Arps ACTL6A/B, the bootstrap support is low (62%, **Fig. 1B, Supplementary Data 1**), suggesting that ACTL8 is too diverged to confidently assign an ancestral gene, whether it was actin or Arp1-10.

The most striking feature of the ACTL8 subfamily is its long branch lengths between orthologs, relative to the very short branches of most canonical Arps (**Fig. 1B**), indicating that ACTL8 diverges more rapidly in sequence across species than the stringently conserved canonical Arps. To determine if residue divergence among ACTL8 orthologs is concentrated within one region of the protein, we generated a predicted structure of ACTL8 using AlphaFold 3^51^ and mapped conservation of ACTL8 orthologs on the surface. Like most Arps, ACTL8 is predicted to exhibit the canonical actin fold, including four subdomains that bind ATP^3^ (**Fig. 1C, Supplementary Fig. 1B**). We observed some patches of residue divergence, most notably in subdomain 3. In contrast, the “pointed” and “barbed” ends of the actin fold, which mediate protein interactions with some Arps and longitudinal interactions within actin filaments^52^, are largely conserved among orthologs (**Supplementary Fig. 1B**). We then analyzed how human ACTL8 diverges with respect to human actin (ACTB). ACTL8 is 34% identical to actin with divergence found throughout the sequence (**Supplementary Fig. 1C**). Furthermore, while ACTL8 is approximately the same length as actin, it exhibits small insertions (1-6 residues) and deletions (1-7 residues) in subdomains 1, 2 and 4 (**Fig. 1C, Supplementary Fig. 1C**). This divergence in tertiary structure from actin is similar to canonical Arps1-3 and contrasts with that of the nuclear Arps, which have large insertions throughout the actin fold^20^.

We compared the tissue expression profile of ACTL8 to that of all canonical Arps and the non-canonical Arps ACTR1B, ACTR3B, and ACTL6B. In the GTEX expression database^34,53^, ACTL8 appears exclusively expressed in the testis (**Fig. 1D**). In contrast, canonical Arps are ubiquitously expressed in all tissue types, except for the paralog ACTL6B, which is predominantly expressed in the brain. Recent RNA-seq data indicates that ACTL8 is also highly expressed in oocytes^54,55^ (**Fig. 1D**). Given its restricted expression profile, ACTL8 may play roles in germ cell development and potentially the early embryo.

Many reproductive proteins evolve under positive selection, such that amino acid sequence changes that occur at a rate far greater than expected under neutral evolutionary drift^56–58^. Given that the ACTL8 subfamily has strikingly long branch lengths and is enriched in expression in the germline (**Fig. 1B, D**), we tested whether ACTL8 evolves under positive selection using primate sequences (see methods). We found that ACTL8 shows statistically significant evidence for positive selection using the PAML methodology^59^ (**Supplementary Fig. 1D**). However, when we masked CpG dinucleotides, which tend to be hypermutable^60,61^, ACTL8 no longer shows a signature of positive selection (**Supplementary Fig. 1D**). Therefore, rapid evolution of ACTL8 could be due to positive selection or to the presence of CpG dinucleotides. Regardless, the rate of sequence changes in ACTL8 strikingly contrasts with that of actin, which evolves under negative (purifying) selection^62,63^. Overall, our evolutionary and expression analyses suggest that ACTL8 is a rapidly evolving, non-canonical Arp that recently arose in mammals with a unique germline-specific expression profile.

### ACTL8 localizes predominantly to the nucleus of spermatogonia and cancer cells upon misexpression

Subcellular localization is closely linked to the diverse functions of the canonical Arps. Thus, we next examined the localization of ACTL8 to gain insight into its molecular role. Members of the Arp superfamily are generally enriched either in the cytoplasm (e.g., actin, Arps 1-3 and Arp10, **Fig. 1B**) or are predominantly in nuclei, where they participate in chromatin remodeling (e.g., Arps 4-8, **Fig. 1B**). We first examined ACTL8 localization in the human testis by immunostaining with an anti-ACTL8 antibody. ACTL8 was detectable in the early stages of sperm development, which are characteristically in the basal compartment of a seminiferous tubule (**Fig. 2A, B**). Strikingly, ACTL8 predominantly colocalized with DNA, with only weak cytoplasmic staining, suggesting it is largely a nuclear Arp in the testis (**Fig. 2B**). We surveyed ACTL8 for a nuclear-localization sequence (NLS), but prediction tools failed to identify any obvious NLS sequence (cNLS mapper^64^, NucPred^65^), suggesting ACTL8 may be imported into the nucleus via another protein. Similarly, actin lacks an NLS and requires Importin-9 for nuclear localization^66–68^.

**Figure 2:**
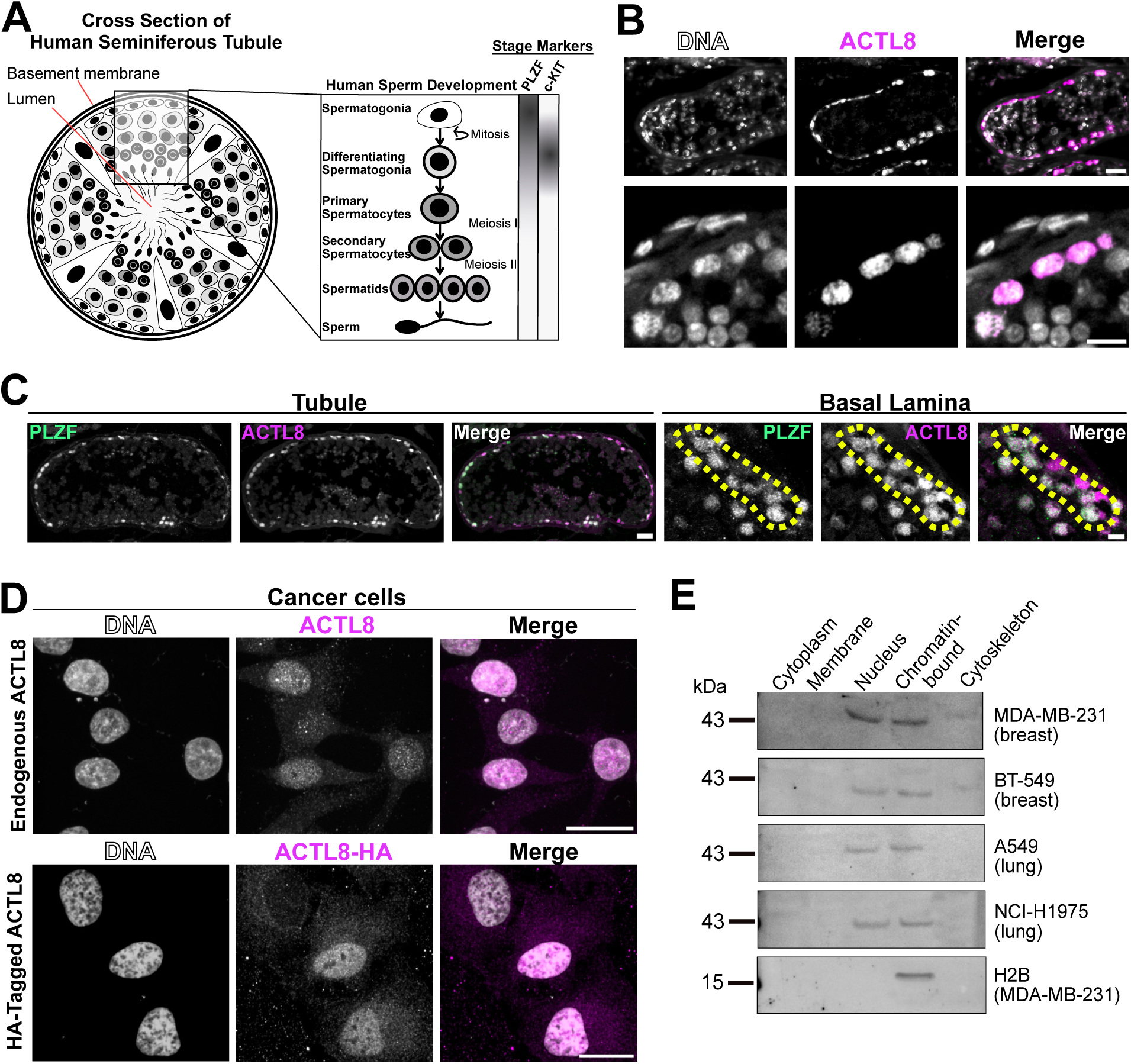
ACTL8 localizes predominantly to the nucleus of spermatogonia and cancer cells upon misexpression. **A)** Schematic cross-section of a human seminiferous tubule illustrating the stages of sperm development from germ cells to mature spermatozoa. Expression patterns of early spermatogenesis markers are indicated as a gradient (darker color reflects higher expression) based on published datasets^132,53,133^. **B)** Images of human testis tissue immunostained for endogenous ACTL8 using an anti-ACTL8 antibody. ACTL8 signal is enriched in nuclei of cells at the periphery of seminiferous tubules. Scale bar: 30 µm. Bottom: higher-magnification view, scale bar: 20 µm. **C)** Co-immunostaining of ACTL8 with PLZF (undifferentiated spermatogonia). Left: whole seminiferous tubule, scale bar: 30 µm. Right: higher magnification of the basal compartment, scale bar: 10 µm. **D)** Images of ACTL8 localization in cancer cells. Top: MDA-MB-231 cells immunostained for endogenous ACTL8 (same antibody as in panel B). Bottom: HeLa cells exogenously expressing ACTL8-HA and stained for the HA tag. Scale bar: 20 µm. **E)** Immunoblots of subcellular fractionations of several cancer cell lines: triple-negative breast cancer (MDA-MB-231, BT-549) and non-small cell lung cancer (A549, NCI-H1975). Probed with anti-ACTL8 antibody and anti-Histone H2B antibody as a chromatin-bound control.

Single-cell RNA-sequencing data of human testes suggests that ACTL8 is first expressed in undifferentiated spermatogonia and then most highly in differentiated spermatogonia^35,55^ (**Supplementary Fig. 2A**). Spermatogonia are at a critical point in spermatogenesis in which transcriptional networks and signaling cascades change, promoting mitosis and differentiation to meiotic spermatocytes^69,70^ (**Fig. 2A**). To precisely identify the stage at which ACTL8 protein is expressed during sperm development, we co-immunostained human testis sections with antibodies against established protein markers for the stages of early sperm development. ACTL8 frequently overlapped with PLZF, a transcription factor expressed in undifferentiated spermatogonia (also known as ZBTB16^71,72^, **Fig. 2C, Supplementary Fig. 2C**), and rarely colocalized with c-KIT (also known as KIT^73–77^, **Supplementary Fig. 2B**), a receptor tyrosine kinase protein-KIT expressed in differentiating spermatogonia. Thus, while human testis RNA-seq data suggests highest transcripts levels in differentiating spermatogonia (**Supplementary Fig. 2A**), ACTL8 protein is found primarily in undifferentiated, PLZF-positive spermatogonia (**Fig. 2C, Supplementary Fig. 2C**).

Because *ACTL8* is aberrantly expressed in several cancer types, we next asked whether its nuclear localization is preserved outside the testis when misexpressed in other cell types. We stained triple-negative breast cancer cells MDA-MB-231, which appear to misexpress *ACTL8* mRNA, using the same commercial anti-ACTL8 antibody and confirmed ACTL8 localizes predominantly in the nucleus (**Fig. 2D, Supplementary Fig. 2D**). We also exogenously expressed HA-tagged ACTL8 in HeLa cells and found that ACTL8-HA, stained with an anti-HA antibody, displayed the same nuclear localization (**Fig. 2D, Supplementary Fig. 2D**). To further validate nuclear localization, we performed subcellular fractionation followed by western blotting of breast and lung cancer cell lines that misexpress *ACTL8*. We found that ACTL8 is enriched in the nuclear and chromatin-bound fractions in all cancer cell lines tested (**Fig. 2E**). Collectively, our findings demonstrate that ACTL8 is a novel nuclear Arp and suggests it likely plays roles in spermatogonia while maintaining its nuclear localization upon misexpression.

### ACTL8 regulates genes involved in cell migration and proliferation

The nuclear roles of actin and canonical Arps ultimately influence gene expression^78^. Thus, we first investigated how ACTL8 impacts gene expression in cancer. We performed RNA sequencing using MDA-MB-231 cells, an epithelial breast cancer cell line, because they express *ACTL8* highly and ACTL8 expression correlates with poorer survival of breast cancer patients^79^ (**Supplementary Fig. 3A**). We compared RNA-seq data between control MDA-MB-231 cells, treated with a “scrambled” siRNA oligo that does not target *ACTL8*, and MDA-MB-231 cells with reduced *ACTL8* expression upon siRNA knockdown (KD) (**Fig. 3A**). We further validated knockdown of *ACTL8* by confirming that exogenously expressed ACTL8-HA is reduced by the siRNA oligos (**Supplementary Fig. 3B**), and each of the four siRNAs efficiently reduced ACTL8 protein levels when tested independently (**Supplementary Fig. 3C**). Our RNA-sequencing data show that ACTL8 knockdown causes widespread transcriptional changes. Using thresholds of a false discovery rate-adjusted p-value ≤ 0.05 and a fold change ≥ 1.5 in mRNA levels (**Fig. 3B, Supplementary Data 5**), we find that 743 genes increase in transcript abundance and 593 decrease. Expression changes for a panel of these genes were validated by quantitative real-time PCR (RT-qPCR) (**Fig. 3C, Supplementary Fig. 3D**). We found with Gene Ontology^80–82^ (GO) enrichment analysis that pathways upregulated upon *ACTL8* reduction were primarily related to cell migration (e.g., positive regulation to wounding), cell proliferation (e.g., nuclear division, chromosome segregation), and cell differentiation (e.g., positive regulation of glial cell differentiation) (**Fig. 3D**). Pathways associated with immune and inflammatory responses (e.g., antigen processing and presentation, MHC protein complex assembly) were also downregulated (**Fig. 3D**).

**Figure 3:**
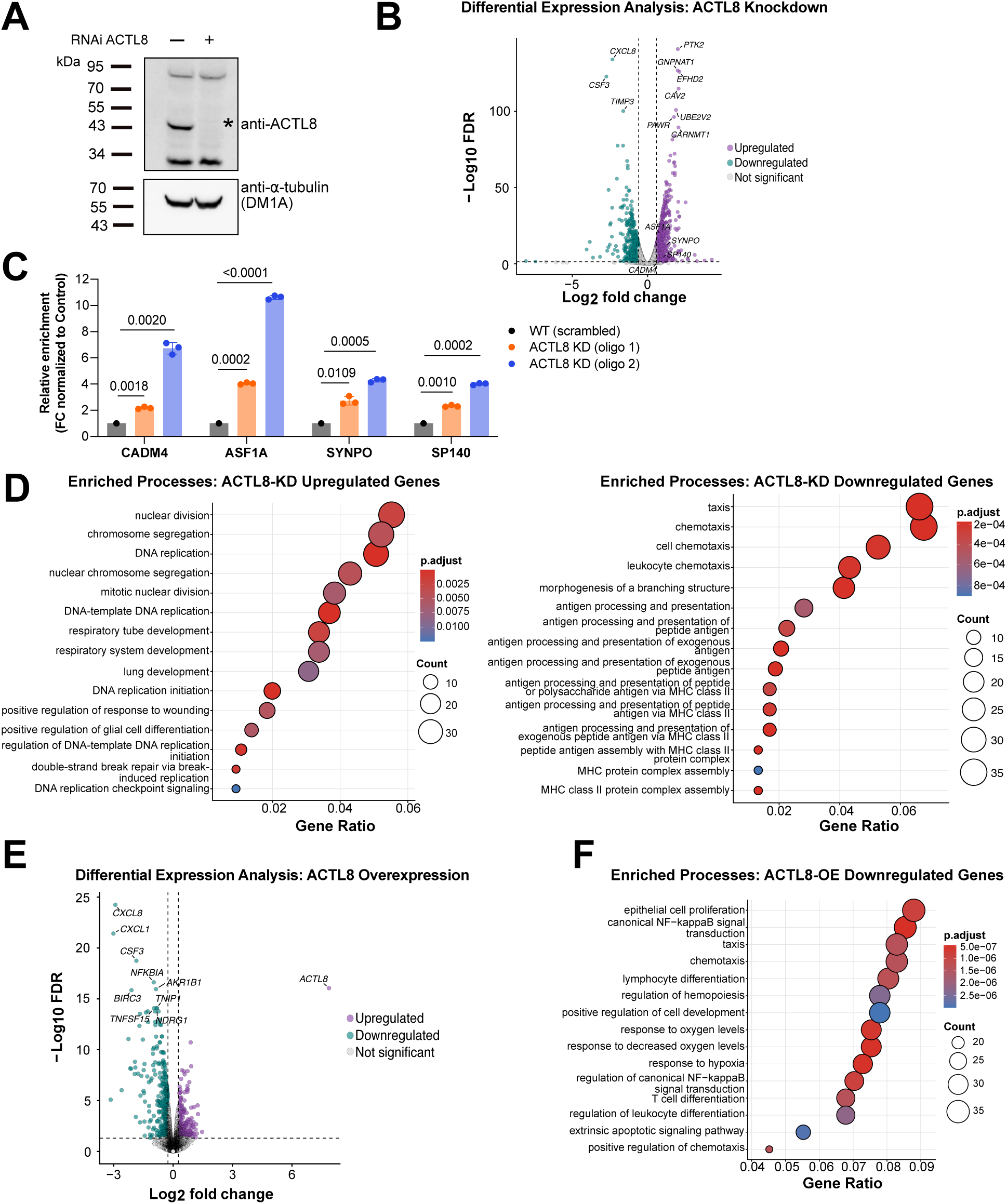
ACTL8 regulates genes involved in cell migration and proliferation. **A)** Immunoblot showing efficient knockdown of *ACTL8* in MDA-MB-231 cells transfected with control (scrambled) siRNA or *ACTL8*-targeting siRNA oligo (pool of four siRNAs targeting *ACTL8* mRNA). ACTL8 is indicated by an asterisk (*), and *a*-tubulin was used as a loading control. **B)** Volcano plot from RNA-seq analysis of *ACTL8* knockdown in MDA-MB-231 cells using the siRNA pool shown in (A), highlighting differentially expressed genes with false discovery rate (FDR) ≤ 0.05 and fold change (FC) ≥ 1.5. Downregulated genes are shown in teal, upregulated genes in purple, and non-significant (NS) genes in grey. Selected genes (top 10 that are most significant and genes analyzed in C) are highlighted with white circles. **C)** RT–qPCR validation of selected genes from the RNA-seq dataset (technical replicates shown). See two additional biological replicates in Fig. S3. Expression levels were normalized to 18S rRNA across samples, and the relative expression of oligo 1- and oligo 2-treated samples was normalized to the WT (scrambled siRNA) condition. Two siRNA oligos were used separately to rule out oligo-specific effects. Oligo 1 targets the coding sequence, and oligo 2 targets the 3’ UTR of ACTL8 mRNA. Statistical analysis: unpaired t-test with Welch’s correction. **D)** Gene Ontology^80–82^ (GO, Biological Process) enrichment analysis of genes upregulated (left) and downregulated (right) upon *ACTL8* knockdown. Gene ratio refers to the ratio between input genes that are associated with a GO term and the total number of input genes that were mapped to the GO database. **E)** Volcano plot from RNA-seq analysis of *ACTL8* overexpression in MDA-MB-231 cells following adenoviral infection, with FDR ≤ 0.05 and FC ≥ 1.2. We lowered the FC threshold to 1.2 from 1.5, as used in the knockdown, to identify more genes impacted by *ACTL8* overexpression, given overexpressing *ACTL8* had less of an impact than the *ACTL8* knockdown. Downregulated genes are shown in teal, upregulated genes in purple, and non-significant (NS) genes in grey. **F)** GO enrichment analysis of genes downregulated upon *ACTL8* overexpression.

To confirm our findings, we took a complementary approach and overexpressed *ACTL8* to test whether increased *ACTL8* expression similarly impacts transcription. We overexpressed *ACTL8* using an adenoviral system that allows for induced *ACTL8* expression via doxycycline (**Supplementary Fig. 3E, F**). We infected MDA-MB-231 cells with adenovirus and conducted RNA-sequencing to compare uninduced versus induced *ACTL8* expression. Using thresholds of a false discovery rate-adjusted p-value ≤ 0.05 and a fold change ≥ 1.2 in mRNA levels, we find that 280 genes increase in transcript abundance and 431 decrease upon ACTL8 overexpression (**Fig. 3E, Supplementary Data 5**). Too few genes were upregulated to perform GO enrichment analysis; thus, we performed this analysis only on downregulated genes and found that ACTL8 overexpression impacted similar pathways to those altered upon ACTL8 knockdown. Processes related to cell movement (e.g. taxis, chemotaxis), cell proliferation (e.g., epithelial cell proliferation), and cell development were transcriptionally downregulated upon *ACTL8* overexpression (**Fig. 3F**). Together, our RNA-sequencing analyses with knockdown and overexpression experiments indicate that *ACTL8* in breast cancer cells predominantly regulates the expression of genes in cell migration and proliferation pathways.

### ACTL8 modulates cell migration

Because ACTL8 influences the expression of genes involved in migration, we tested whether reducing ACTL8 does in fact alter migration using a wound-healing assay in MDA-MB-231 (triple-negative breast cancer) and HCC-1954 (HER2+ breast cancer) cell lines. Following 48 hours of siRNA treatment, we scratched confluent cell monolayers to create a wound and monitored its closure over time with live-cell imaging (**Fig. 4A**). We found that *ACTL8*-depleted cells closed the wound faster than control cells in both MDA-MB-231 (*p*=0.0412, 0.0338, **Fig. 4B, C, Supplementary Movies 1 and 2**) and HCC-1954 cells (*p*=0.0494, 0.0355 **Fig. 4D, Supplementary Fig. 4A**). Given this result with siRNA-mediated knockdown, we expected that overexpression of *ACTL8* should have the opposite effect—reduced migration. Using our inducible *ACTL8* construct, we compared induced versus uninduced *ACTL8* expression in MDA-MB-231 cells and found that increased *ACTL8* expression does indeed slow migration (*p*=0.0442, **Fig. 4A, E**). The magnitude of the effect when overexpressing ACTL8 is less than that of the knockdown. This difference may be due to MDA-MB-231 cells already expressing ACTL8 and a maximum rate of wound closure is reached at a certain point of expression. Collectively, these results demonstrate that ACTL8 slows migration of breast cancer cells, aligning with our transcriptomic analyses that indicate ACTL8 alters the expression of gene programs controlling migration, either directly or indirectly.

**Figure 4:**
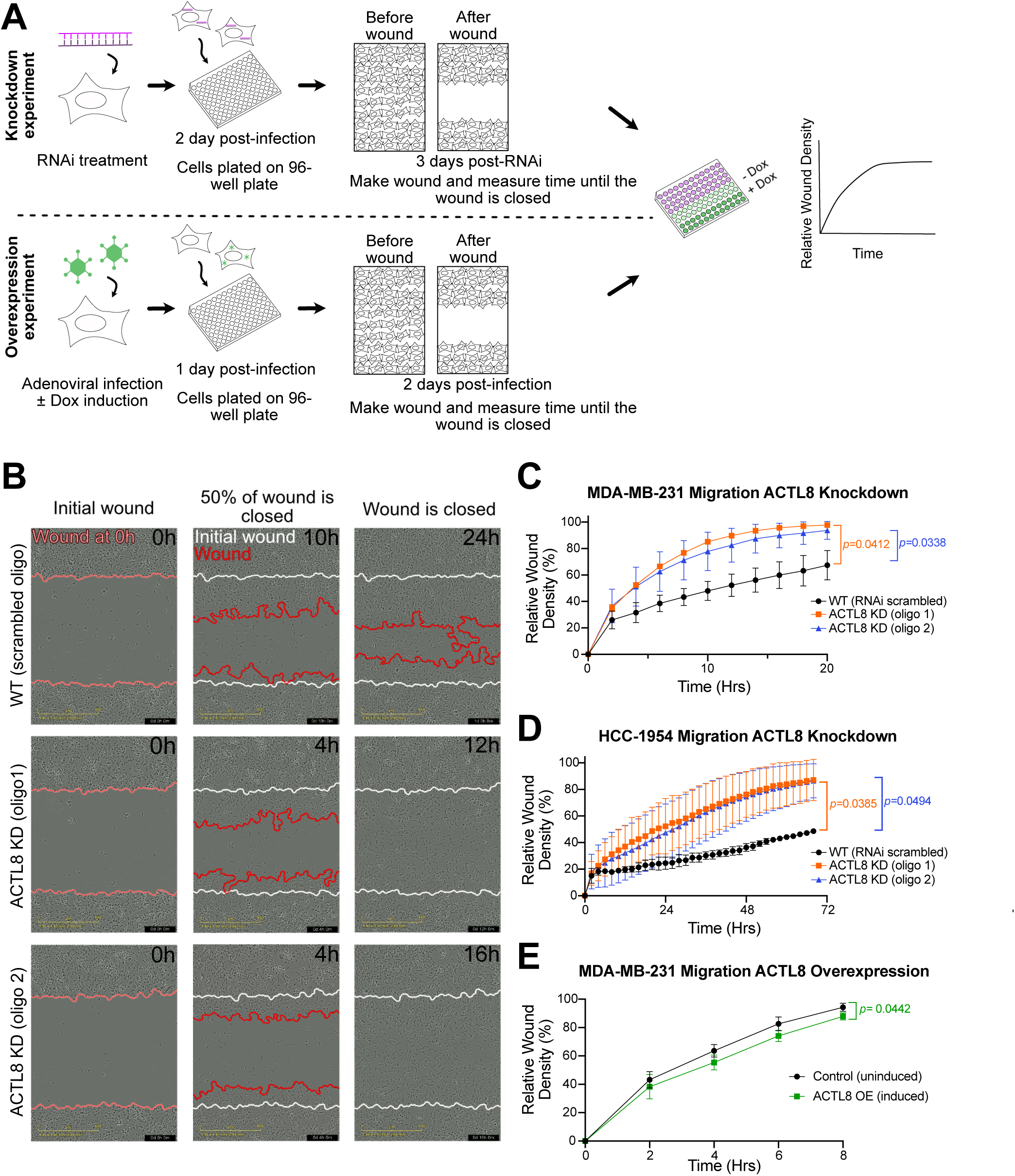
ACTL8 modulates cell migration. **A)** Schematic of the workflow for migration assays. Cells were treated with siRNAs or with doxycycline for 2 days prior to seeding in 96-well plates. Wounds were generated at 3 days post-RNAi treatment or 2 days post-infection, and wound closure was monitored over time, with images acquired every two hours until ∼95-98% wound closure. **B)** Representative images of time points during wound-healing assays with MDA-MB-231 cells, transfected with scrambled siRNA (control) or *ACTL8*-targeting siRNAs (“oligo 1” and “oligo 2”). Images show the initial wound (0 h), ∼50% wound closure, and complete wound closure. Time required to reach each stage is indicated in the top right of each image. Wound boundaries were automatically marked by the Incucyte SX5 scratch wound analysis software module. **C-D)** Quantification of migration shown as relative wound density over time in MDA-MB-231 or HCC-1954 cells treated with scrambled siRNA (black) or individual *ACTL8*-targeting siRNAs (orange and blue). Data represent three independent biological replicates, each performed with three technical replicates. **E)** Quantification of migration shown as relative wound density over time in MDA-MB-231 upon induced or uninduced *ACTL8* overexpression. Data represent three independent biological replicates, each performed with three technical replicates. Statistical significance was assessed at the final time point using an unpaired t-test with Welch’s correction in C-E.

Because genes involved in cell proliferation were transcriptionally altered in our RNA-sequencing data, we also tested whether reduced expression of *ACTL8* affects cell count following siRNA treatment. Although *ACTL8* knockdown showed a trend toward increased proliferation in both MDA-MB-231 and HCC-1954 cell lines, the effect was not as robust as found with cell migration and lacked statistical significance due to high variability across replicates (**Supplementary Fig. 4B**).

### ACTL8 interacts with the histone lysine demethylase KDM5B

To gain mechanistic insight into how ACTL8 may alter transcription and migration, we sought to identify putative protein interactors of ACTL8. We performed a yeast two-hybrid (Y2H) screen using a human testis-enriched cDNA library to identify interactors where ACTL8 is natively expressed. This screen yielded many hits that are nuclear proteins, but one in particular, the histone lysine demethylase KDM5B (also known as PLU-1 or JARID1B)^83^, drew our interest due to its role in chromatin biology. KDM5B catalyzes the removal of mono-, di- and trimethyl groups from lysine 4 of histone H3 (H3K4me1/2/3), a chromatin mark associated with transcriptional activation^81^. Due to its demethylation of H3K4, KDM5B is considered to normally act as a transcriptional repressor^48^. Like ACTL8, KDM5B is predominantly expressed in the testis^35,55^ (**Supplementary Fig. 5A**) and misexpressed in many cancers^85^. As expected^86^, we found that KDM5B is expressed in our MDA-MB-231 cell line, even when ACTL8 is knocked down (**Supplementary Fig. 5B**) and is predominantly nuclear (**Supplementary Fig. 5C**). Therefore, while KDM5B was identified using a testis cDNA library, it is also a likely interactor in cancer.

We further validated the interaction between ACTL8 and KDM5B by performing co-immunoprecipitation (co-IP) experiments with exogenously expressed HA-tagged ACTL8 and V5-tagged KDM5B in HEK-293T cells. We found that KDM5B-V5 bound the bait ACTL8-HA with minimal non-specific binding to anti-HA resin (**Fig. 5A**). We next sought to map the minimal region of KDM5B mediating the interaction with ACTL8. KDM5B contains multiple structural domains: a catalytic JmjN/JmjC domain, an AT-rich, DNA-binding domain (ARID), a zinc finger domain (ZnF), and three plant homeodomain domains (PHD1, 2 and 3)^86,87^ (**Fig. 5B**). In the Y2H screen, ACTL8 interacted with a fragment of KDM5B that included the PHD2 domain and two flanking α-helices— one upstream and one downstream of PHD2 (**Supplementary Fig. 6A, B**). PHD fingers generally serve as protein-interacting domains. Among the KDM5B PHD fingers, PHD1 binds unmethylated H3K4 (H3K4me0), and PHD3 shows highest affinity for H3K4me3, whereas PHD2 appears to lack histone-binding activity^86^. By co-IP, we found that the Y2H fragment including the PHD2 domain robustly interacted with ACTL8 (**Fig. 5C**). We further tested whether the PHD2 domain alone, excluding the flanking *a*-helices, was sufficient for the interaction, but we failed to detect any binding (**Supplementary Fig. 6C**). The individual PHD1 and PHD3 domains also showed no binding (**Supplementary Fig. 6D, E**). We next tested whether the upstream α-helix fused to PHD2 (“α1-PHD2”) or PHD2 fused to the downstream α-helix (“PHD2-α2”) was sufficient for binding ACTL8. PHD2-α2 robustly interacted with ACTL8, while α1-PHD2 failed to bind (**Fig. 5D, Supplementary Fig. 6F**). Given that PHD2 alone was not sufficient for the interaction (**Supplementary Fig. 6C**), we next tested whether the downstream *a*-helix alone could bind ACTL8. The “α2” helix alone did interact with ACTL8 (**Supplementary Fig. 6G**), albeit weakly compared to the PHD2-α2 fragment (**Supplementary Fig. 6H**). All together, these data demonstrate the binding interface between ACTL8 and KDM5B includes the *a*-helix immediately downstream of the PHD2 domain, while the PHD2 likely strengthens this interaction. Because this region of KDM5B does not directly bind histones^86^, it is possible that the ACTL8 interaction with KDM5B does not block histone binding.

**Figure 5.**
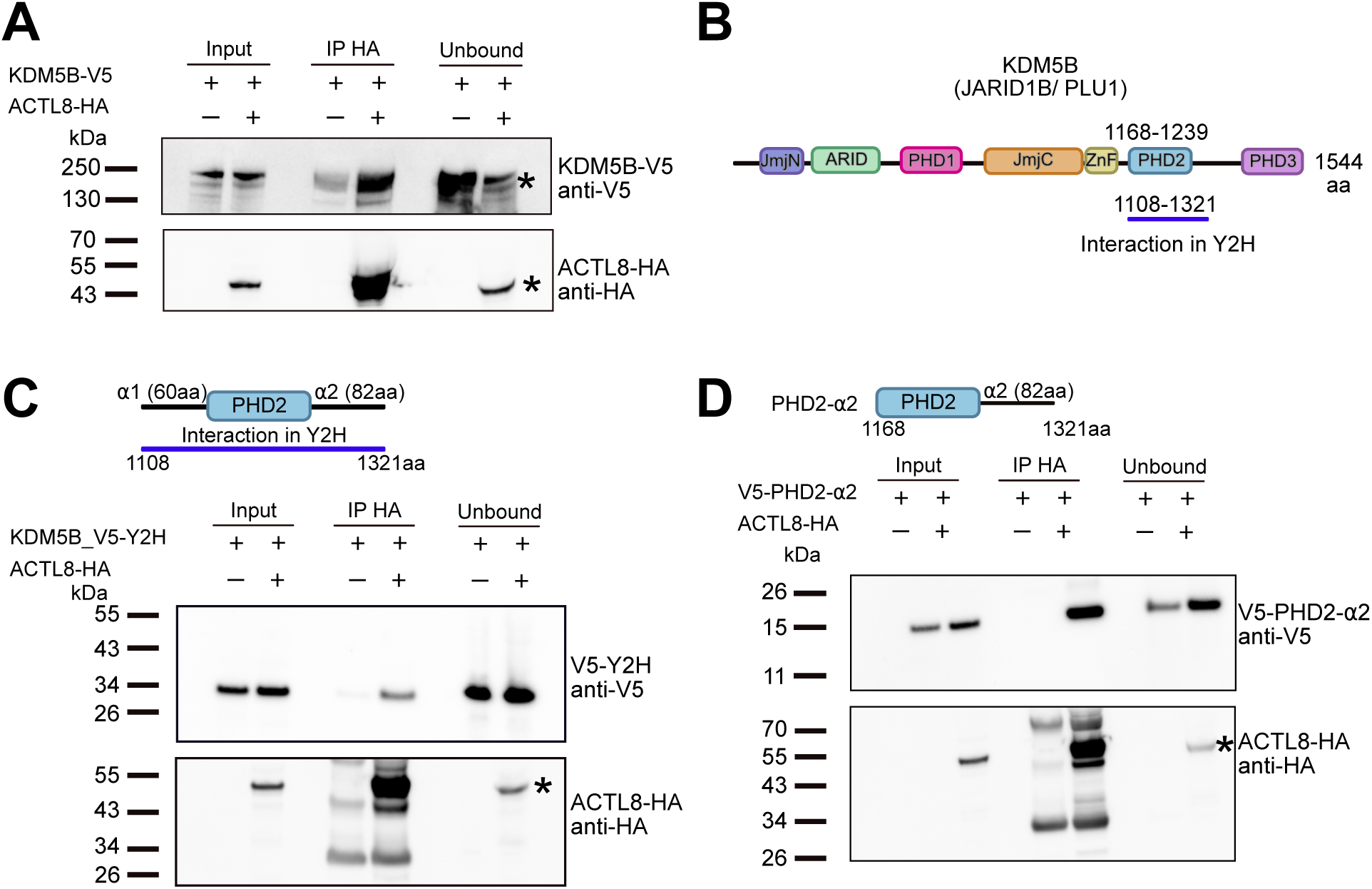
ACTL8 interacts with the histone lysine demethylase KDM5B. **A)** Immunoblot of ACTL8-HA and KDM5B-V5 co-immunoprecipitation (co-IP). HEK-293T cells expressing KDM5B-V5 with and without ACTL8-HA were incubated with anti-HA magnetic beads and immunoblotted for HA and V5 tags. Asterisks indicate bands corresponding to ACTL8 or KDM5B. **B)** Schematic of KDM5B domain structure (amino acid positions were assigned based on Klein et al., 2014^86^), highlighting the ACTL8-interacting region identified in the yeast two-hybrid (Y2H) screen with corresponding residues noted. **C)** Immunoblot of co-IP as in panel A with HEK-293T cells expressing ACTL8-HA and a fragment of KDM5B-V5, the ACTL8-interacting region identified by the Y2H screen. Asterisk indicates ACTL8. **D)** Immunoblot of co-IP as in panel A with HEK-293T cells expressing ACTL8-HA and the V5-tagged PHD2-*a*2 fragment of KDM5B. Asterisk indicates ACTL8.

### Loss of ACTL8 increases H3K4me3 predominantly at genes that regulate migration

Next, we investigated how ACTL8 influences KDM5B function. Given that KDM5B is an H3K4 demethylase, we mapped chromatin occupancy of H3K4me3, an active histone mark typically enriched at gene promoters and transcription start sites (TSS)^88^, genome-wide by CUT&RUN^89^ to test whether its levels or genomic distribution are altered upon ACTL8 reduction. In parallel, we mapped H3K27me3 as a control for a repressive histone mark^90^. We performed CUT&RUN with two different breast cancer cell lines that express *ACTL8*, MDA-MB-231 and HCC-1954, the same cell lines used in the migration assays (**Fig. 4**). We compared control cells (treated with a scrambled siRNA oligo) to cells treated with *ACTL8*-targeting siRNA oligos. Knockdown efficiency was confirmed by western blotting for each replicate (**Supplementary Fig. 7A**), and the two biological replicates from both cell lines showed high reproducibility in signal across the genome (**Supplementary Fig. 7B**). We first examined how H3K4me3 changed at genes that increased in expression in MDA-MB-231 cells upon ACTL8 depletion (**Fig. 3B**). Given that H3K4me3 is associated with active transcription, we would expect upregulated genes in our previous RNA-seq analyses to gain H3K4me3. Indeed, we found that, on average, both replicates with MDA-MB-231 cells exhibited an increase in H3K4me3 occupancy proximal to the TSSs of upregulated genes (**Fig. 6A**). We next asked how H3K4me3 changed globally in both MDA-MB-231 and HCC-154 cells and observed that loss of ACTL8 resulted in increased H3K4me3 by at least 1.4 fold in a significant number of peaks in both cell lines, albeit fewer in MDA-MB-231 cells (**Fig. 6B**). As done with our RNA-sequencing data, we performed a GO enrichment analysis of genes associated with peaks that gained H3K4me3 signal upon *ACTL8* knockdown and found that in both cell lines, increases in H3K4me3 signal were primarily associated with genes that impact cell migration (**Fig. 6C, D**), consistent with the transcriptome changes in migration-related pathways observed in our RNA-sequencing analyses (**Fig. 3D**). Processes that involve large changes in migration, including blood vessel development and angiogenesis, were also significantly represented (**Fig. 6C, D**). We verified that the distribution of signal in H3K4me3-enriched regions was largely consistent in the control and knockdown samples between replicates for both cell lines, with far more peaks exhibiting an increase in H3K4me3 upon ACTL8 loss (**Fig. 6E**). In contrast, H3K27me3 changes were modest (**Supplementary Fig. 8**).

**Figure 6:**
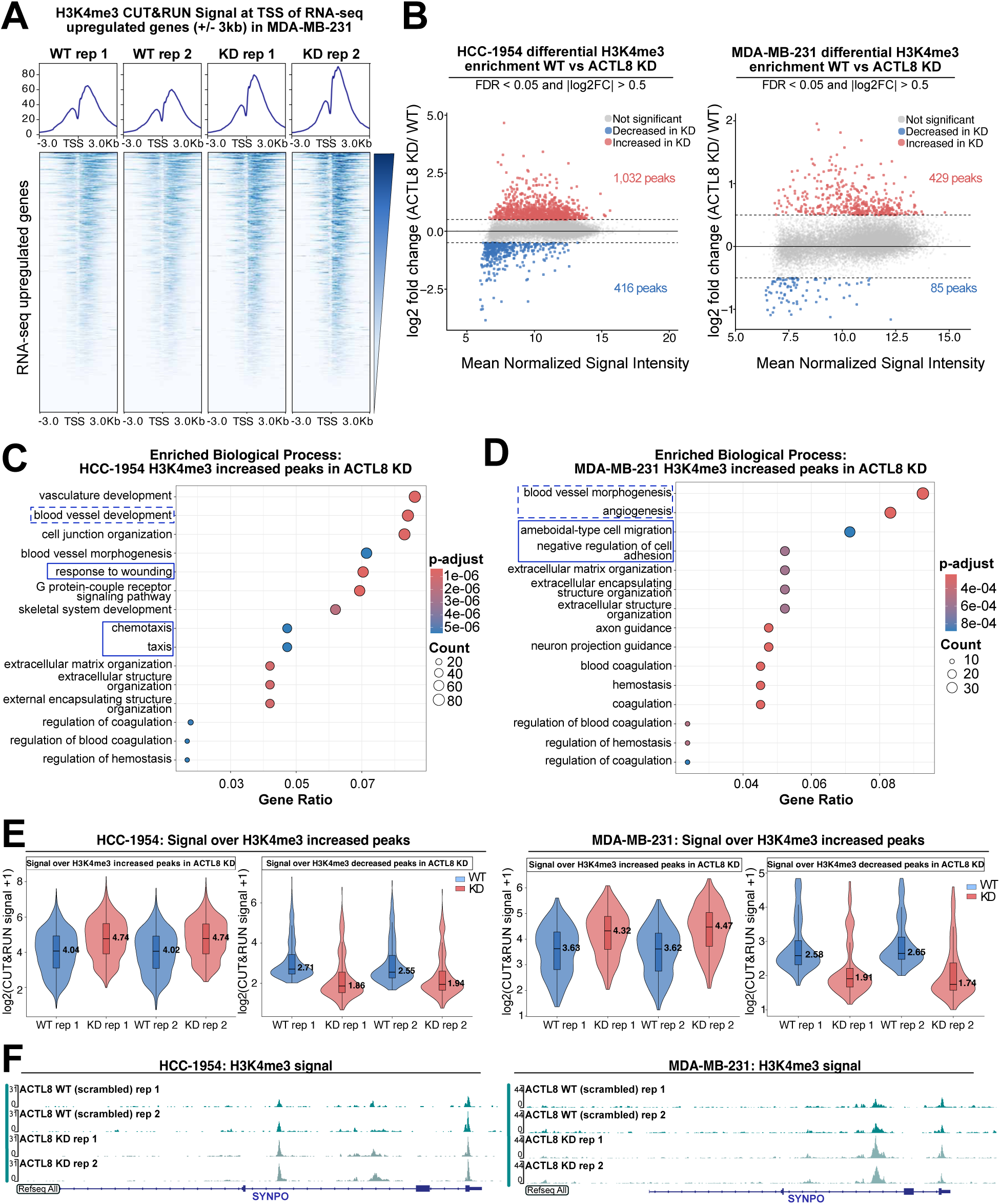
Loss of *ACTL8* alters H3K4me3 predominantly at genes regulating migration. **A)** Heatmaps and average profile plots showing H3K4me3 CUT&RUN signal centered on the transcription start sites (TSS) (±3 kb) of genes identified as upregulated in the *ACTL8* knockdown RNA-seq dataset (Fig. 3B). Signal from *ACTL8* WT (scrambled siRNA) and *ACTL8* knockdown (KD) is shown for two biological replicates with MDA-MB-231 cells. Regions are ordered by descending H3K4me3 signal intensity, and average profiles are displayed above the corresponding heatmaps. **B**) MA plots showing differential H3K4me3 enrichment between *ACTL8* WT (scrambled siRNA) and *ACTL8* knockdown (KD) in HCC-1954 (left) and MDA-MB-231 (right) cells. Peaks were identified by DiffBind^117^, and differential enrichment was determined using a false discovery rate (FDR) < 0.05. Each point represents an H3K4me3 peak plotted by mean normalized signal versus log₂ fold change (KD/WT). Red and blue points denote peaks with increased or decreased H3K4me3 in *ACTL8* knockdown (KD), respectively, with an absolute log₂ fold change > 0.5, while grey points indicate peaks with lesser or non-significant differences. **C-D**) Gene Ontology (Biological Process) enrichment analysis of H3K4me3 peaks increased in *ACTL8* KD in HCC-1954 (C) and MDA-MB-231 (D) cells. Gene Ratio is shown on the x-axis, dot size represents gene count, and color indicates adjusted p-value (“p.adjust”). GO terms associated with cell migration are outlined with solid blue rectangles, while dotted blue rectangles indicate GO terms that may also be associated to cell migration and/or cancer. **E)** To validate differential H3K4me3 regions identified by DiffBind, violin plots showing H3K4me3 CUT&RUN signal over *ACTL8* KD-increased and *ACTL8* KD-decreased peaks were plotted for HCC-1954 (left) and MDA-MB-231 (right) cells. KD-decreased regions exhibit higher H3K4me3 signal in WT samples than in *ACTL8* KD samples, whereas *ACTL8* KD-increased regions show higher signal in KD samples, validating the differential peak classification. Signal is shown as log₂ (normalized CUT&RUN read count + 1) for two biological replicates, with boxes indicating the interquartile range and median values annotated within each violin. **F)** IGV tracks of a representative locus (*SYNPO*), showing increased H3K4me3 occupancy upon *ACTL8* KD in HCC-1954 and MDA-MB-231 cells.

We next examined representative loci with increased H3K4me3 occupancy following *ACTL8* knockdown. In both HCC-1954 and MDA-MB-231 cell lines and across replicates, *SYNPO*, coding for an actin-associated protein involved in cell shape and motility^91^, displayed increased H3K4me3 with decreased H3K27me3, consistent with a more transcriptionally active chromatin state (**Fig. 6F, Supplementary Fig. 9A**). Not all H3K4me3 changes were at the same genes between cell lines. For example, in MDA-MB-231 cells, *CADM4*, which is involved in cell-cell adhesion^92^, displayed increased H3K4me3 with a modest decrease in H3K27me3 (**Supplementary Fig. 9B**); consistent with this change, *CADM4* was upregulated in our RNA-seq data and RT-qPCRs in MDA-MB-231 cells, as was *SYNPO* (**Fig. 3B, C, Supplementary Fig. 3D**). However, *CADM4* showed modest changes in H3K4me3 in HCC-1954 cells (**Supplementary Fig. 9B**). Although the chromatin changes sometimes differed between the two cell lines at a subset of genes, migration-related pathways were enriched in both HCC-1954 and MDA-MB-231 cells (**Fig. 6C, D**), suggesting that ACTL8 influences similar biological processes regardless of which genes are impacted. Together, these data support a model in which ACTL8 functions with KDM5B to alter the genomic landscape at a subset of loci and modulate cell migration.

### The ACTL8-KDM5B interaction contributes to the impact on cell migration

To investigate whether the ACTL8-KDM5B interaction underlies the previously observed migration phenotype (**Fig. 4**), we leveraged the minimal ACTL8-binding region of KDM5B that displayed robust binding, the PHD2-α2 fragment (**Fig. 5D**), as a competitive inhibitor. Because the PHD2-α2 fragment interacts with ACTL8, we hypothesized that overexpressing PHD2-α2 would sequester endogenous ACTL8 from the catalytically active, full-length KDM5B, thereby mimicking *ACTL8* loss. We overexpressed PHD2-α2 in MDA-MB-231 cells using the same doxycycline-inducible system used for *ACTL8* overexpression (**Fig. 3**, **Fig. 4**) and performed a wound-healing assay (**Fig. 7A**). We compared induced versus uninduced PHD2-α2 expression and found that PHD2-α2 expression accelerated cell migration (*p*= 0.0003, **Fig. 7B, C, Supplementary Movies 3 and 4**), similar to *ACTL8* knockdowns in MDA-MB-231 cells (**Fig. 4B, C**). These findings demonstrate that the ACTL8-KDM5B interaction contributes to the regulation of cell migration and suggest it ultimately reduces migration, likely through transcriptional repression of relevant pathways observed in our genomic analyses (**Fig. 6C, D**).

**Figure 7:**
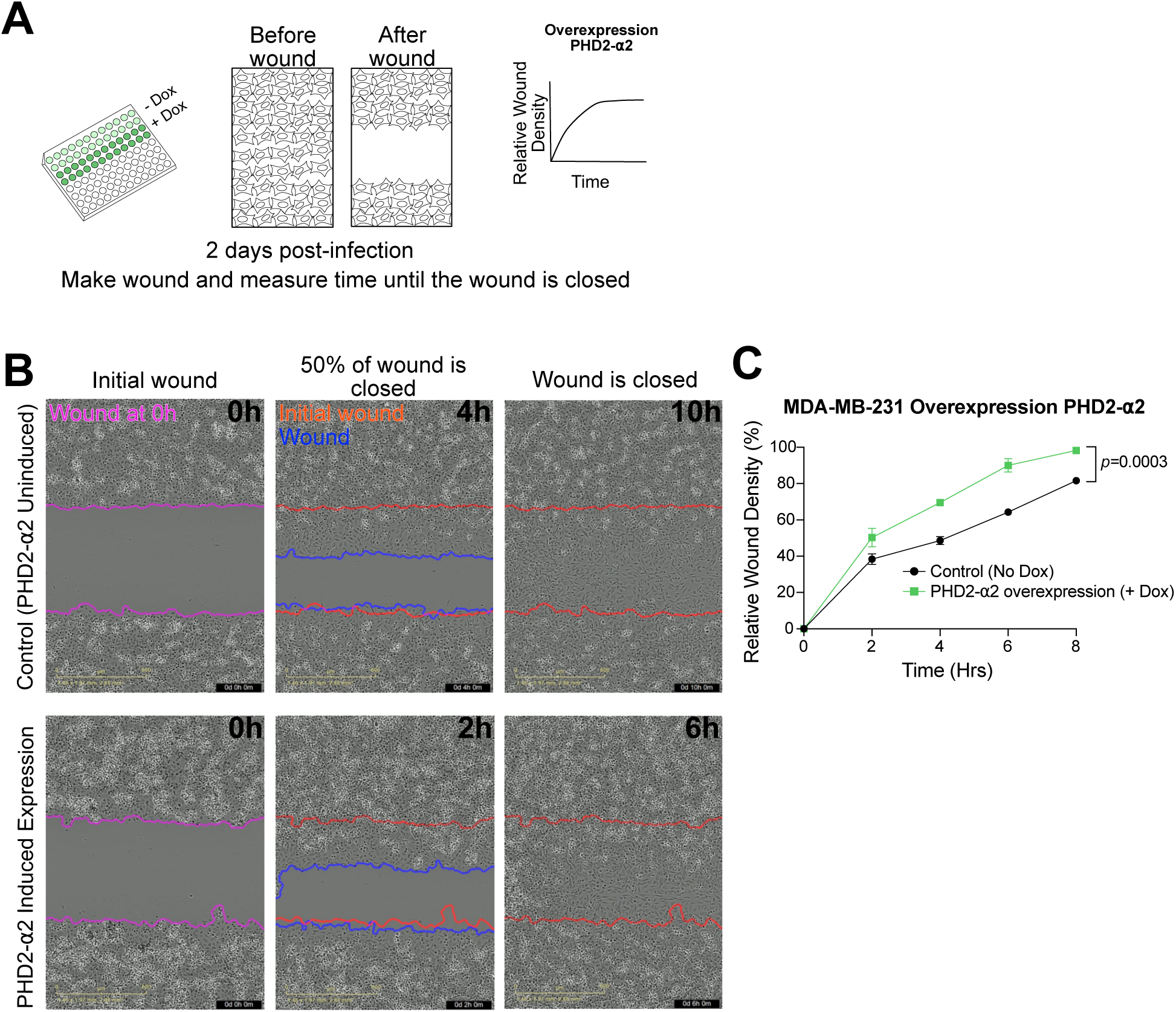
The ACTL8-KDM5B interaction contributes to the impact on cell migration. **A)** Schematic of the workflow for migration assays. After a 4 h adenoviral infection, PHD2-*a*2 expression was induced or uninduced with doxycycline (“Dox”) for 2 days. Cells were then plated on a 96-well plate, wounds were generated, and wound closure was monitored over time until closure, acquiring images every two hours. **B)** Representative images of time points during wound-healing assays with MDA-MB-231 and PHD2-*a*2 overexpression uninduced (no Dox) or induced (plus Dox). Images show the initial wound (0 h), ∼50% wound closure, and complete wound closure. The time required to reach each stage is indicated in the top right of each image. **C)** Quantification of migration shown as relative wound density over time, with or without PHD2-*a*2 expression. Data represent three independent biological replicates, each performed with three technical replicates. Statistical significance was assessed at the final time point using an unpaired t-test with Welch’s correction.

## Discussion

Unlike the ubiquitously expressed canonical members of the Arp superfamily, *ACTL8* exhibits germline-specific expression with uncharacterized molecular functions. Here, our findings reveal the evolutionary history, nuclear localization, and molecular role of ACTL8 in chromatin biology. We demonstrate that in breast cancer cells, ACTL8 alters the expression of a subset of genes, consequently reducing cell migration. We further show that ACTL8 interacts with the histone H3 lysine demethylase KDM5B, a transcriptional repressor, and this interaction contributes to ACTL8-mediated changes in cell migration. Upon *ACTL8* loss, H3K4me3 levels increase at specific loci involved in cell migration, suggesting that ACTL8 supports KDM5B activity, either directly or indirectly, at a subset of genes. Overall, our study identifies ACTL8 as a unique nuclear Arp that plays a role in KDM5B-mediated transcriptional repression, ultimately regulating migratory behavior.

Similar to ACTL8, KDM5B is enriched in the testis and ovary and, being upregulated in cancer, is also classified as a cancer/testis antigen^93^. Interestingly, KDM5B displays a phenotype similar to that of ACTL8 in the triple-negative breast cancer cell line MDA-MB-231; it suppresses cell migration and appears to play a tumor-suppressive role^86^. However, the effect of KDM5B expression appears to be cell-type dependent. For example, in ER+ breast cancer cells, KDM5B has been reported to promote, rather than suppress, cell proliferation and invasion^48,94,95^. This phenotypic contrast may also reflect different expression levels, as KDM5B is expressed more highly in ER+ breast cancer cells than in triple-negative breast cancer cells^86^. Thus, KDM5B can promote neoplastic behavior or act as a tumor suppressor depending on the context^96^. A similar context-dependent role may apply to ACTL8, given its functional parallels with KDM5B. It will be important in future studies to determine whether ACTL8 promotes or suppresses oncogenic phenotypes across different cancer contexts, including ER+ breast cancer. Indeed, previous reports have described that ACTL8 may promote cell motility, invasion, and proliferation in other cancer types^28,41–45,47,97^. These findings contrast with our observation of a negative effect on migration and may be due to cell-type-specific effects or differences in expression levels, similar to what has been reported for KDM5B. Moreover, while it is tempting to link the suppressive migratory behavior of ACTL8 we observed to cancer progression in patients, migration speed into a wound does not always positively correlate with the aggressiveness of a cancer^98^. Regardless of how ACTL8 ultimately impacts cancer progression, our findings demonstrate that ACTL8 affects cancer cell biology by modulating specific transcriptional programs.

We find that the PHD2 finger and a neighboring *a*-helix in KDM5B contain the minimal binding interface for ACTL8. Interestingly, the PHD2 finger appears to be the least conserved among the PHD fingers across KDM5 family members (KDM5A–D, ∼40% pairwise identity versus 50% for PHD1 and 60% for PHD3, **Supplementary Fig. 10**). The lower sequence similarity of PHD2 is even more pronounced compared to the highly conserved catalytic domain across KDM5s (>80%). Like the PHD2 finger, the upstream *a*-helix (α2) also shows low sequence similarity across KDM5s (∼32% identity). We hypothesize that the divergence of the PHD2–*a*2 region allows for KDM5B-specific interactions, such as with ACTL8. Because only PHD1 and PHD3 bind histones, PHD2-interacting proteins like ACTL8 may not perturb histone binding but rather regulate KDM5B activity at histones. Based on our findings that H3K4me3 increased at specific loci upon ACTL8 loss, ACTL8 may facilitate either the recruitment of KDM5B to specific target genes or modulate its activity at these sites. Future work will distinguish between these two possibilities.

ACTL8 may perform roles that other nuclear Arps fail to carry out, including regulating KDM5B activity. The evolutionary history of the Arp superfamily suggests that the functionally versatile actin fold has been recurrently co-opted for specialized roles, and the restricted expression of ACTL8 suggests it acquired a specialized germline role in mammals. Spermatogonia, in which ACTL8 is natively expressed, are a unique cell type found only in the testis and may require a tissue-specific Arp like ACTL8 to modulate their distinct transcriptomes prior to differentiation and meiosis^99,100^. Furthermore, undifferentiated progenitor spermatogonia exhibit critical migratory behavior in the testis^101,102^, which ACTL8 may regulate, as it does in breast cancer cells. KDM5B is also expressed during early sperm development, where it has been proposed to regulate the transcription of genes involved in meiotic progression^103^. Although it remains technically challenging to dissect directly the function of ACTL8 and its interaction with KDM5B in the human germline, emerging systems that allow for studying human sperm development *ex vivo*^104^ may enable identification of how ACTL8 regulates the chromatin landscape in the mammalian germline.

The signatures of rapid evolution found in ACTL8 (**Supplementary Fig. 1D**) further suggest that its function is evolving across species, unlike the stringently conserved canonical Arps. The testis often exhibits elevated levels of rapidly evolving proteins due to strong selective pressures from sexual selection and sperm competition^56,57^. ACTL8 may help shape species-specific chromatin biology in early sperm development. Overall, our work demonstrates that ACTL8 is a unique example of an evolutionarily divergent mammalian Arp deployed for cell-type-specific chromatin regulation.

## Methods

### Cell Lines

MDA-MB-231, NCI-H1975, BT-549 and A549 cell lines were obtained from Dr. Angelique Whitehurst (UT Southwestern). HCC-1954 cells were obtained from Dr. Srinivas Malladi (UT Southwestern). HEK-293T cell line was obtained from Dr. Samir M. Parikh (UT Southwestern). HeLa cells were obtained from Dr. Lee Kraus (UT Southwestern). Breast cancer cells MDA-MB-231s were maintained in high glucose DMEM (Sigma-Aldrich #D6429) supplemented with 10% FBS (Sigma-Aldrich #12306C). NCI-H1975s were maintained in RPMI-1640 (Sigma-Aldrich # R8758) supplemented with 5% FBS, and HCC-1954s were maintained in RPMI-1640 supplemented with 10% FBS, 1X penicillin/streptomycin (Cytiva #SV30010), 2mM glutamax (Gibco #35050061) and 1 µg/ml amphotericin B (Gibco #15290018). BT-549s were maintained in RPMI-1640 supplemented with 5% FBS and A549s were maintained in RPMI-1640 supplemented with 5% FBS. HEK-239T cells were maintained in high glucose DMEM supplemented with 10% FBS and 25 mM HEPES pH 7.4. HeLa cells were maintained in in high glucose DMEM (Sigma-Aldrich #D6429) supplemented with 10% FBS (Sigma-Aldrich #12306C). Adeno-X 293 cells (Takara #632271) were maintained in high glucose DMEM supplemented with 10% FBS, 1X penicillin/streptomycin and 4 mM Glutamax according to Takara Adeno-X Adenoviral system 3 user manual. All the cell lines were maintained at 37°C with 5% CO2 and split every 2-3 days. Cells were tested for mycoplasma contamination with a PCR detection kit (SouthernBiotech #130001). Authenticity of MDA-MB-231, NCI-H1975, BT549, HCC-1954 was verified by extracting genomic DNA using a DNeasy blood and tissue kit (Qiagen #13323) and genotyped with the PowerPlex® Fusion System for detection of 13 core loci in the Combined DNA Index System (CODIS) (done by the McDermott Center Sanger Sequencing Core at UT Southwestern). To passage or harvest cells, cells were washed with 1X PBS (Quality Biological #25-507) and trypsinized with Trypsin-EDTA (Sigma-Aldrich #T3924). The reaction was quenched by adding complete growth medium (cell line–specific medium supplemented with FBS as described above).

### Testis Histology

#### ACTL8 and PLZF immunostaining

Formalin-fixed paraffin embedded testis sections (Tissue Array.com #HuFPT151) were baked for 1 h at 60°C. Sections were treated with 3 washes of xylene (mixed isomers), followed by 2 washes of 100% ethanol, and 1 wash each of 95%, 80%, and 70% ethanol for 5 min each. Antigen retrieval was performed at 90-95°C for 20 min in 10 mM Sodium Citrate Buffer (pH 6) for sections co-stained for PLZF and ACTL8. Sections were cooled in tap water for 10 min, permeabilized in 1X PBS containing 0.1% Triton X-100 (PBST) for 30 min, and blocked in blocking solution for 2 h at room temperature. Blocking solution for sections immunostained for ACTL8 was 3% BSA and 5% Animal Serum in PBST. Blocking solution for sections co-immunostained for ACTL8 and PLZF was 5% BSA in PBS. Sections were incubated overnight at 4°C with primary antibodies against ACTL8 (Abcam #ab184562; 1:100) or PLZF (R&D Systems #AF2944; 1:500) diluted in blocking solution. The following day, sections were washed three times with PBST for 5 min each, incubated for 2 h at room temperature with fluorescently conjugated secondary antibodies (anti-rabbit 647, Invitrogen #A-21245; anti-rabbit 488 Invitrogen #A-11034; anti-mouse 488, Invitrogen #A-21235; anti-goat 647 Invitrogen #A-21447) all diluted 1:200 in blocking solution, and washed three times with 1X PBS for 5 min each. All antibodies used are listed in **Supplementary Table 1**. Nuclei were stained with 1:5000 diluted Hoechst (Invitrogen #H3570) in 1X PBS, followed by two 1 min washes in MilliQ water. Sections were allowed to air-dry and mounted using ProLong Diamond Antifade Mountant (Invitrogen #P36965).

#### c-KIT immunostaining

Testis samples were fixed in 4% paraformaldehyde (PFA) overnight at 4°C, washed in 1X PBS, incubated in 30% sucrose for 24 h, and frozen in O.C.T. Five µm sections were prepared and light exposed for 4-6 h to reduce autofluorescence prior to blocking. Slides were incubated for 45 min at room temperature in blocking solution (PBS containing 3% BSA and 0.1% Triton X-100) and subsequently incubated with primary antibody against c-KIT (R&D Systems #AF1350; 1:1000) or ACTL8 (Abcam #ab184562; 1:500) diluted in blocking solution for 1 h at room temperature. Sections were incubated 15 min in wash solution (PBS containing 0.1% Triton X-100). Fluorescently conjugated secondary antibodies (anti-goat 555 Invitrogen #A-21432 and anti-rabbit 488 Invitrogen #A-21206) all diluted 1:500 in blocking solution were added and incubated in blocking solution at room temperature for 1 h, followed by 15 min in wash solution. All antibodies used are listed in **Supplementary Table 1.** Coverslips were applied over sections with Vectashield containing DAPI (Vector Laboratories #H-1200). Negative controls were conducted by omitting primary antibodies.

### siRNA-Mediated Knockdown

siRNAs were reverse-transfected using Lipofectamine RNAiMAX transfection reagent (Invitrogen #13778075) according to the manufacturer’s instructions. Briefly, siRNAs were diluted in Opti-MEM I Reduced Serum Medium, no phenol red media (Gibco #11058021) to a final concentration of 50 nM for RNAi experiments with pooled siRNA oligos targeting ACTL8 or 100 nM for single siRNA experiments (siRNAs oligo 1, 2, 3 and 4). Lipofectamine RNAiMAX was then added to the diluted siRNA at a 1:1 ratio, and the siRNA–lipid complexes were dispensed into 6-well plates or 10-cm^2^ dishes. After a 20 min incubation at room temperature, resuspended cells were added directly to the siRNA-lipid mix. Complete growth medium was subsequently added to reach a final volume of 2 ml (6-well plate) or 10 ml (10 cm^2^ dish). The MISSION siRNA Universal Negative Control #1 (Sigma-Aldrich # SIC001) was used as negative control (RNAi scrambled). The siRNAs against ACTL8 were purchased from Dharmacon, ON-TARGETplus Human ACTL8 siRNA SMARTPool Actl8 (Dharmacon #L-014660-02-0005) for RNAi *ACTL8* pool experiments, and for single siRNA experiments siRNAs oligo 1 (Dharmacon #J-014660-17-0005, targeting CDS), oligo 2 (Dharmacon #J-014660-18-0005, targeting 3’ UTR), oligo 3 (Dharmacon #J-014660-19-0005, targeting CDS) and oligo 4 (Dharmacon #J-014660-20-0005, targeting 3’ UTR).

### Migration and Cell Proliferation Assays

To assess cell migration, wound-healing assays were performed with three independent biological replicates. Two days after siRNA treatment or one day after adenoviral infection, 5 x 10^4^ MDA-MB-231 or HCC-1954 cells were seeded into collagen-coated (Cell Applications #125-50) for MDA-MB-23, or Matrigel-coated (Corning #356230) for HCC-1954 96-well plates, with three wells used as technical replicates per biological replicate. Cells were incubated at 37°C with 5% CO2 for 24 h to allow attachment. Uniform scratches were generated using a 96-well WoundMaker (Sartorius # BA-04858). Wells were washed twice with 1X PBS to remove detached cells, and fresh complete growth medium was added. Cell migration into the wound area was monitored using the Incucyte SX5 Live-Cell analysis system (Sartorius #4816) until wound closure (∼8-20 h for MDA-MB-231 cells and ∼68-72 h for HCC-1954, see imaging details under “Microscopy and Image Analysis”).

To assess cell proliferation, 7.5 x 10^4^ MDA-MB-231 or HCC-1954 cells were siRNA-treated for two days and then seeded onto coverslips in 6-well plates. Proliferation assays were performed with three independent biological replicates. Cells were incubated at 37°C with 5% CO2 for four days, cells attached to the coverslips were washed once with cold 1X PBS and fixed for 15 min with 4% PFA (prepared by diluting 20% PFA (Electron Microscopy Sciences #50-980-492) in 1X PBS). Fixed cells were washed three times for 5 min each with 1X PBS, 0.2% Triton X-100 (PBST). Nuclei were stained with 1:1000 diluted Hoechst (Invitrogen #H3570) in 1X PBS for 1 min, followed by one wash with 1X PBS. Coverslips were allowed to air-dry and mounted onto glass slides using ProLong Diamond Antifade Mountant (Invitrogen #P36965). Thirteen images per sample from three independent biological replicates were acquired (see “Microscopy and Image Analysis”).

### Immunofluorescence

For endogenous ACTL8 localization assays, MDA-MB-231 cells were seeded onto coverslips in 6-well plates and incubated at 37°C with 5% CO2 for 24 h to allow attachment. For exogenous ACTL8 localization assays, HeLa cells were transfected using FuGENE HD transfection reagent (Promega #E2691) at a 3:2 reagent-to-DNA ratio (see Transfections and immunoprecipitations for details) and fixed three days post-transfection for 15 min with 4% paraformaldehyde (PFA). Fixed cells were washed three times for 5 min each with 1X PBS containing 0.2% Triton X-100 (PBST) and then blocked in 3% bovine serum albumin (BSA; Sigma-Aldrich #A4503) diluted in 1X PBS, for 2-3 h at 4°C. Cells incubated overnight at 4°C with primary antibodies against ACTL8 (Abcam #ab184562) or anti-HA (Invitrogen #26183) diluted in 3% BSA. The following day, cells were washed three times for 5 min each with PBST and incubated for 1 h at room temperature with Alexa Fluor 647–conjugated secondary antibodies (anti-rabbit, Invitrogen #A-21245 or anti-mouse, Invitrogen #A-21235) diluted in 3% BSA and protected from light. Following three washes, nuclei were stained with Hoechst (1:1000 dilution in 3% BSA; Invitrogen #H3570) for 1 min, followed by one final wash with 1X PBS. All antibodies used are listed in **Supplementary Table 1**. Coverslips were mounted using ProLong Diamond Antifade Mountant (Invitrogen #P36965).

### RNA-Sequencing and Analyses

For RNA-sequencing, RNA was extracted from three independent biological replicates. siRNA-mediated knockdown in MDA-MB-231 cells was performed as described above including control samples (RNAi scrambled) in 6-well plates. Three days after siRNA treatment, cells were washed once with 1X PBS. After removal of PBS, 1 ml of TRIzol reagent (Invitrogen #15596026) was added to each well, and cells were lysed by repeated pipetting up and down. Lysates were transferred to microcentrifuge tubes, and 250 µl of chloroform was added per 1 ml of TRIzol. Samples were vortexed for 15 s, incubated at room temperature for 5 min, and centrifuged at 10,000 rpm for 10 min at 4°C. After centrifugation, the top layer (aqueous layer) was transferred to a fresh microcentrifuge tube, mixed with 550 µl of isopropanol, and incubated at room temperature for 5 min. RNA was pelleted by centrifugation at 14,000 rpm for 30 min at 4°C. Supernatant was carefully removed, and RNA pellet was washed with 1 ml of 75% ethanol diluted in OmniPur DEPC-treated water (Sigma-Aldrich #9602-500). Samples were centrifuged at 9,500 rpm for 5 min at 4°C, ethanol was removed, and RNA pellets were air-dried before resuspension in 22 µl of OmniPur DEPC-treated water. RNA concentration was measured using a NanoDrop spectrophotometer (Thermo Fisher Scientific ND-ONE-W). Prior to RNA cleanup, 8 µg of total RNA per sample was treated with DNase I (Zymo #E1010) at a final concentration of 0.2 U/µl for 1 h at 37°C. Following DNase I treatment, RNA was purified using the RNA Clean & Concentrator kit (Zymo #R1014) according to the manufacturer’s instructions. RNA was eluted in 10 µl OmniPur DEPC-treated water (Sigma-Aldrich #9602-500).

*ACTL8 Knockdown experiments:*

#### RNA Quality Assessment and Library Preparation

For RNA-sequencing of ACTL8-knockdown experiments, samples were submitted to the Genomics Core at UT Southwestern. The Agilent 2100 Bioanalyzer system and the RNA Nano kit (Agilent Technologies #5067-1511) were used for RNA quality measurement. All 12 samples had RNA Integrity Number (RIN) scores of 10. The Illumina TruSeq Stranded mRNA Library Preparation Kit (Illumina #20020594) was used for library construction. The total RNA input for each library ranged from 2–3 µg per sample. The workflow included purification of poly(A)+ mRNA using oligo-dT magnetic beads, mRNA fragmentation, first- and second-strand cDNA synthesis, end repair and A-tailing, ligation of UMI adapters (manufactured by Integrated DNA Technologies), followed by two rounds of AMPure XP bead purification, and PCR amplification (9 cycles) to generate the final libraries. Library quantity and quality were then assessed. Concentration was measured using the Quant-iT™ PicoGreen dsDNA Assay Kit (Invitrogen #P7589) on a PerkinElmer Victor X3 (2030 Multilabel Reader). Library quality was evaluated using an Agilent 2100 Bioanalyzer with the DNA High Sensitivity Kit (Agilent Technologies #5067-4626). All 12 libraries met QC requirements prior to sequencing.

#### Library Sequencing and Data Analysis

The samples were sequenced on an Illumina NovaSeq 6000 platform using a paired-end 2 × 150 bp protocol. After demultiplexing, data quality metrics were generated and reviewed. Paired end demultiplexed fastq files were generated using bcl2fastq2 (Illumina, v2.20), from NovaSeq 6000 bcl files. Initial quality control was performed using FastQC v0.11.8 and multiqc v1.7. Fastq files were imported batch wise, trimmed for adapter sequences followed by quality trimming using QIAGEN CLC Genomics Workbench (CLC Bio, v24.0.2, https://digitalinsights.qiagen.com). The imported high-quality reads were mapped against gene regions and transcripts annotated by ENSEMBL v104 GRCh39 using the RNA-Seq Analysis tool v2.6 (QIAGEB, CLC Genomics Workbench), only matches to the reverse strand of the genes were accepted (using the strand-specific reverse option). Differential gene expression analysis between the sample groups was done using the Differential Expression for RNA-seq tool v2.7, where the normalization method was set as TMM. Differential expression between the groups was tested using the control group, outliers were downweighed, and filter on average expression for FDR correction was enabled.

#### ACTL8 Overexpression

For expression changes upon ACTL8 overexpression, total RNA was extracted as described above and submitted to the company Plasmidsaurus for RNA-sequencing. Library preparation, sequencing using Illumina technology, and primary data processing (including read alignment and quantification) were performed by the provider using standard pipelines. Custom analysis and annotation were also carried out by Plasmidsaurus (https://plasmidsaurus.com/technical-documentation/rna).

### Differential Expression Analyses and RNA-seq Visualization

Gene Ontology (GO)^80–82^ enrichment analysis was performed in R (v4.5.1) using the clusterProfiler package^105^. Enrichment was carried out using the Biological Process ontology with Benjamini–Hochberg correction, using all annotated human genes as “the background universe”. Terms with an adjusted p-value and q-value below 0.05 were considered significant. Results were visualized as dot plots showing the top 15 enriched terms ranked by adjusted p-value, with dot size representing gene count and dot color reflecting statistical significance.

Volcano plots were generated in R (v4.5.1) using the ggplot2^106^ and ggrepel^107^ packages. Protein-coding genes were classified as significantly upregulated or downregulated if they met a fold change threshold of ≥ 1.5 (or ≥ 1.2 where indicated) and an FDR-adjusted p-value ≤ 0.05. For ACTL8-knockdown RNA-seq datasets, an additional max group mean cutoff of > 2 was applied. Each gene is plotted by its log₂ fold change against −log₁₀(FDR).

### Real-Time Quantitative PCRs

RNA from three independent biological replicates was extracted and purified as for RNA-sequencing. Following RNA clean-up, cDNA was synthesized using the SuperScript III First-Strand Synthesis System for RT-PCR (Invitrogen #18080-051) according to manufacturer’s instructions. Briefly, 8 µg of purified RNA was reverse-transcribed using oligo(dT) primers (50 µM) to generate cDNA. For each sample, a no-reverse-transcriptase control was included in which the reverse transcriptase enzyme was replaced with water. RT-qPCRs were performed using iTaq Universal SYBR Green Supermix (BioRad #1725121) with approximately 50 ng of cDNA per reaction on an Applied Biosystems 7500 Real-Time PCR System (Life Technologies), following the manufacturer’s recommendations. Amplification was carried out using a standard curve method with the following cycling conditions: 95°C for 15 sec and 60°C for 60 sec, for a total of 40 cycles. Primers were picked based on single peaks in the melt curve analyses, and their specificity was evaluated by using cDNA from wild-type cells as the template. Primer sequences are listed in **Supplementary Table 2**^108–110^. 18S rRNA served as a “housekeeping gene” to normalize cDNA quantity across samples. Three technical replicates were performed for each biological replicate to control for pipetting variability.

### Immunoblots

Trypsinized cells were collected, transferred to conical tubes, and centrifuged at 0.5 x g for 5 min. Supernatants were removed, and cell pellets were resuspended in RIPA buffer (Teknova #R3792) supplemented with 1X cOmplete EDTA-free protease inhibitor cocktail (Roche #11836170001) using 25 µl per well of a 6-well plate or 300 µl per 10-cm^2^ dish. Samples were transferred to microcentrifuge tubes and incubated on ice for 50 min to allow cell lysis. Samples were then centrifuged at maximum speed for 10 min at 4°C, and the supernatants containing soluble protein were transferred to fresh tubes. NuPAGE LDS Sample Buffer (Invitrogen #NP0007) and NuPAGE Sample Reducing Agent (Invitrogen #NP0009) were added to each lysate to a final concentration of 1X. Samples were boiled at 95°C for 5 min and run on NuPAGE Bis-Tris 4–12% Mini Protein Gels (Invitrogen #NP0322BOX) using 1X NuPAGE MES SDS Running Buffer (Invitrogen #NP0002) at 190 V for 35 min. Proteins were transferred onto PVDF membranes using iBlot 2 Transfer Stacks (Invitrogen #IB24002) and the iBlot 2 Dry Blotting System (Invitrogen #IB21001) with the P0 program (20 V for 1 min, 23 V for 4 min, and 25 V for the remainder of a 7 -min run) or 20V for 9 min when probing for anti-KDM5B. Membranes were blocked for 20 min at room temperature in 5% milk in Tris-buffered saline (Sigma-Aldrich #T5912) with 0.1% Tween-20 (TBST). Membranes were incubated overnight at 4°C with primary antibodies (listed in **Supplementary Table 1**), washed three times for 5 min each with TBST, and then incubated with the appropriate secondary antibodies (listed in **Supplementary Table 1**) for 1 h at room temperature. Following three additional washes with TBST, proteins were visualized using Clarity Western ECL Substrate (Bio-Rad #1705060) or by fluorescence scanning at 647 nm using Image Lab software (Bio-Rad).

### Transfections and Co-Immunoprecipitations

One day prior to transient transfections, 0.9 x 10^6^ HEK-293T cells were seeded into 10 cm^2^ dishes, with one dish used per transfection. Transfections were performed using FuGENE HD transfection reagent (Promega #E2691) at a 3:1 reagent-to-DNA ratio, diluted in OptiMEM I Reduced Serum Medium, no phenol red media (Gibco #11058021). For co-transfections, 1 µg of each plasmid DNA was used (2 µg total DNA) in a final volume of 600 µl of FuGENE-DNA-Opti-MEM mixture, whereas for single-plasmid transfections, 1 µg of DNA was used. Three days post-transfection, cells were harvested as described above and lysed in 300 µl of lysis buffer (50 mM Tris-HCl pH 7.4, 150 mM NaCl, 1% NP-40 and 5 mM EDTA) supplemented with 1x cOmplete EDTA-free protease inhibitor cocktail (Roche #11836170001) and 1 mg/ml BSA (Sigma-Aldrich #A4503). Samples were transferred to microcentrifuge tubes and incubated on ice for 50 min to allow cell lysis. Samples were then centrifuged at maximum speed for 10 min at 4°C, and the supernatants containing soluble protein were transferred to fresh tubes. An aliquot of 5 µl of each lysate was reserved as “input”. The remaining lysate was incubated with 25 µl of pre-washed anti-HA magnetic beads slurry (Thermo Fisher Scientific # 88837) for 1 h at room temperature with rotation. Beads were pre-washed three times with TBS with 0.05% Tween-20. Following incubation, beads were collected using a magnetic stand and the unbound fraction was retained. Beads were washed five times with lysis buffer, and during the final wash, beads were transferred to a fresh microcentrifuge tube before magnetic separation. Immunoprecipitated samples were eluted by adding 25 µl of 1X NuPAGE LDS Sample Buffer (Invitrogen #NP0007) and 1X NuPAGE Sample Reducing Agent (Invitrogen #NP0009).

### Subcellular Fractionation

One 10-cm^2^ dish of cells at ∼80% confluency was used for each cell line. Subcellular fractionation was performed using the Subcellular Protein Fractionation Kit (Thermo Fisher Scientific #78840), following the manufacturer’s instructions. For fractionating only cytoplasm versus nucleus (**Supplementary Fig. 5A**), one 10-cm^2^ dish of MDA-MB-231 cells at ∼80% confluency was used. Subcellular fractionation was performed using a combination of the Subcellular Protein Fractionation Kit (Thermo Fisher Scientific #78840) and the NE-PER Nuclear and Cytoplasmic Extraction Reagents (Thermo Fisher Scientific #78833). The cytoplasmic extract was first prepared according to the manufacturer’s instructions for NE-PER Nuclear and Cytoplasmic Extraction Reagents kit. Briefly, cells were harvested as described above and pelleted by centrifugation at 500 x g for 5 min. The pellet was washed and resuspended in 1X PBS without Ca^2+^, Mg^2+^, Phenol Red (Quality Biological #25-507), followed by centrifugation at 500 x g for 3 min. After removal of the supernatant, the pellet was resuspended in ice-cold Cytoplasmic Extraction Reagent I (CER I) supplemented with 1X cOmplete EDTA-free protease inhibitor cocktail (Roche #11836170001). Samples were vortexed at the highest setting for 15sec and incubated on ice for 10 min. Cytoplasmic Extraction Reagent II (CER II) was then added, followed by vortexing for 5 sec at the highest setting and incubated on ice for 1 min. Samples were vortexed again and centrifuged at maximum speed for 5 min. The supernatant corresponding to the cytoplasmic fraction was transferred to a fresh microcentrifuge tube and retained for further analysis (step 5 in Cytoplasmic and Nuclear Protein Extraction section of the NE-PER Nuclear and Cytoplasmic Extraction Reagents kit, Thermo Fisher Scientific #78833). The remaining pellet was resuspended in NEB buffer supplemented with protease inhibitors, 5 µl of 100 mM CaCl_2_, and 3 µl of Micrococcal Nuclease per 100 µl of NEB to isolate the chromatin-bound fraction (from the Subcellular Protein Fractionation Kit). The samples were vortexed for 15 sec at maximum speed and incubated for 15 min at room temperature. Following incubation, samples were vortexed again for 15 sec and centrifuged at maximum speed for 5 min. The supernatant (chromatin-bound fraction) was transferred to a fresh tube. All fractions were analyzed by western blotting as described above.

### Plasmid Construction

The cDNA encoding human ACTL8 was obtained from UT Southwestern’s McDermott Center for Human Growth and Development, which offers The Ultimate ORF Lite human cDNA collection (Life Technologies). Two common variants of ACTL8 exist according to NCBI’s Single Nucleotide Polymorphism Database^111^ (dbSNP), and they differ by a single amino acid at position 3: alanine (GCA) or serine (TCA). In this study, the variant encoding Ser3 was used. HA-tagged ACTL8 was cloned into the pCMV vector backbone (Addgene plasmid #21039; http://n2t.net/addgene:21039 ; RRID:Addgene_21039) between the AgeI and BamHI restriction sites using the Gibson Assembly system (New England Biolabs #E2611L). The PCR-amplified ACTL8 fragment was purified using the DNA Clean & Concentrator kit (Zymo Research #D4014). The cDNA encoding human KDM5B was obtained in the pDONR222 vector (DNASU) and subcloned into the pLX302 vector (obtained from Dr. Angelique Whitehurst, UT Southwestern) using the Gateway Cloning system (Invitrogen #11791100). Plasmids encoding KDM5B fragments were generated by cloning the corresponding regions into the pWPI vector (from Dr. Angelique Whitehurst, UT Southwestern), linearized with PmeI, using pLX302-KDM5B-V5 as a template and the Gibson Assembly system. Linearized vectors and PCR fragments were gel-purified using the Zymoclean Gel DNA Recovery kit (Zymo Research #D4008), and plasmids were isolated from bacteria using the ZymoPURE Plasmid Miniprep kit (Zymo Research, #D4211). Whole Plasmid Sequencing was performed by Plasmidsaurus using Oxford Nanopore Technology with custom analysis and annotation. Plasmids generated in this study are listed in **Supplementary Table 3**.

### Generation of Recombinant Adenovirus

Adenoviruses driving expression of ACTL8 and the PHD2-*a*2 fragment were generated using the Adeno-X adenoviral system 3 (Takara Bio #631180) according to manufacturer’s instructions. The ACTL8 construct was rendered RNAi-resistant by introducing synonymous substitutions into the coding sequence at the regions targeted by oligos 1 and 3 (see siRNA-Mediated Knockdown), without altering the encoded amino acid sequence. Briefly, ACTL8 and PHD2-*a*2 were cloned into pAdenoX-Tet3G and transformed into Stella competent cells (Takara Bio #636763). Recombinant constructs were linearized with PacI and transfected into Adeno-X 293 cells using the CalPhos Mammalian Transfection Kit (Takara Bio, #631312). Cells were harvested upon cytopathic effect (CPE) (∼7 days post-transfection), lysed, and the lysate used to re-infect Adeno-X 293 cells for viral amplification. Following a second amplification (∼7 days), cells were harvested at CPE and lysed to obtain viral stocks. MDA-MB-231 cells were infected with viral stocks (1:100 dilution). Transgene expression was induced with doxycycline (1 µg/ml) for 48 h, after which cells were either fixed and stained for ACTL8 (**Supplementary Fig. 3E**), harvested for RNA-sequencing (**Fig. 3E**), or used for migration assays (**Fig. 4E and Fig.7**).

### CUT&RUN

#### Sample preparation

CUT&RUN was performed using the CUTANA ChIC/CUT&RUN kit (EpiCypher #14-1048) following the manufacturer’s instructions (Kit version 5, user manual version 5.0) combined with the nuclei extraction protocol for CUTANA assays (EpiCypher). Nuclei Extraction buffer was prepared by supplementing the extraction buffer with Spermidine (EpiCypher #21-1026), protease inhibitor cocktail (EpiCypher #21-1027) and 0.01% digitonin (from 5% Digitonin, EpiCypher #21-1004k). Purified DNA was eluted in 0.1X TE buffer, and DNA concentration was quantified using a Qubit fluorometer (Thermo Fisher Scientific #Q33226) with the Qubit 1X dsDNA Broad Range Assay Kit (Thermo Fisher Scientific #Q33265). Antibodies used for CUT&RUN assays are listed in **Supplementary Table 1**.

#### Library Preparation and Sequencing

For MDA-MB-231 cells, the library preparation and sequencing were performed by Novogene. Libraries were prepared using KAPA Hyper Prep Library Prep kit (Roche) according to the manufacturer’s instructions. Following library amplification, DNA concentration was quantified using Qubit assay (Thermo Fisher Scientific) and library quality was assessed using TapeStation system (Agilent Technologies). Samples were sequenced on an Illumina NovaSeq X Plus 25B instrument with an average depth of 40 million paired-end reads per sample (20 million in each direction), corresponding to around 6Gb per sample.

For HCC-1954 cells, library preparation and sequencing was done by McDermott Center Sanger Sequencing Core at UT Southwestern. CUT&RUN-enriched DNA (∼0.1–5 ng) from HCC-1954 cell line was converted into sequencing libraries using the CUTANA™ CUT&RUN Library Prep Kit Version 1 (EpiCypher #14-1001). DNA fragments were end-repaired, phosphorylated, and 3′ dA-tailed prior to ligation of Illumina-compatible adapters. Following adapter ligation, uracil-specific excision was performed to linearize ligated products, and DNA was purified using magnetic beads. Libraries were then amplified 14 cycles using PCR with dual indexing primers (i5 and i7) to enable multiplexed sequencing. Final libraries were purified, quantified by Qubit fluorometer (Thermo Fisher Scientific), and evaluated for fragment size distribution via capillary electrophoresis on the Agilent Tapestation 4200 (Agilent Technologies). Libraries were normalized, pooled then sequenced on an Illumina NovaseqX with 150×150 paired-end reads, targeting 25 million reads per sample.

#### Data processing

Sequencing data were processed as previously described^112^ (see: https://www.protocols.io/view/cut-amp-tag-data-processing-and-analysis-tutorial-e6nvw93x7gmk/v1). Briefly, adapters were clipped and paired-end reads were mapped to the *Homo sapiens* genome (UCSC Hg38) using Bowtie2^113^ with parameters: --very-sensitive-local --soft-clipped-unmapped-tlen --no-unal --no-mixed --no-discordant --dovetail --phred33 -I 10 -X 1000 --threads 8. Spike-in reads were mapped to the *E. coli* genome with parameters: --end-to-end --very-sensitive --dovetail --no-unal --no-mixed --no-discordant -q --phred33 -I 10 -X 1000 - -threads 6. Continuous-valued data tracks (bedGraph and bigWig) were generated using genomecov in bedtools^114^ v2.30.0 (-bg option) and normalized using genome coverage scaling.

#### Data analysis

Genome-wide correlations between samples were computed using deepTools (v3.5)^115^ . BigWig files were summarized in 1 kb bins using multiBigwigSummary, and Pearson correlations were calculated using plotCorrelation. Peaks were called using SEACR^116^ in the stringent mode with a threshold of 0.01 (top 1% peaks). SEACR uses a signal block approach, which is agnostic to region width, allowing us to call peaks for both H3K4me3 and H3K27me3 using the same parameters. Differential enrichment between wild-type (WT) and knockdown (KD) conditions was assessed using DiffBind^117^. Peak files and corresponding BAM files from two replicates per condition were provided as input. A consensus peak set was generated by retaining peaks detected in at least two of the four samples. Peaks were not recentered or resized thereby preserving the variable widths of broad H3K27me3-enriched regions. Reads from each BAM file were quantified within the consensus peak set, and differential enrichment between KD and WT conditions was assessed using DESeq2^118^. All tested consensus regions were retained for ranked analyses, while significantly differential regions were defined using an FDR (False Discovery Rate) threshold of 0.05 and classified as KD-enriched (referred to in the text as *increased in ACTL8 KD*) or WT-enriched (referred to in the text as *decreased in ACTL8 KD*) using DiffBind log_2_ fold-change cut offs of >0.5 or <-0.5, respectively. Peaks were assigned to their nearest annotated genes using bedtools^114^ closest with a GRCh38 TSS-to-TES gene annotation file. The resulting gene lists were subjected to Gene Ontology enrichment analysis using clusterProfiler::enrichGO^119^ with the Biological Process ontology and Benjamini–Hochberg multiple-testing correction.

#### Data visualization

Heatmaps were generated using deepTools by plotting normalized BigWig signal over BED-defined promoter regions. Promoters corresponding to genes with increased RNA-seq signal following knockdown were extended by 3 kb upstream and downstream, relative to the TSS, prior to matrix generation. Signal over the WT- and KD-enriched peak sets was quantified using multiBigwigSummary from deepTools^115^, which summarizes normalized BigWig signal across genomic regions. Signal distributions were visualized in R using ggplot2^106^. Violin plots with embedded boxplots were used to compare normalized signal across WT and KD conditions and between replicates. Differential enrichment was visualized using MA plots^118^ generated from the DiffBind^117^ results. Regions meeting both the FDR and fold-change thresholds were classified as KD-enriched (FDR < 0.05 and log_2_ fold change > 0.5) or WT-enriched (FDR < 0.05 and log_2_ fold change < −0.5), while all remaining regions were classified as not significant. Horizontal reference lines indicated zero log_2_ fold change and the log_2_ fold-change thresholds of ±0.5.

### Microscopy and Image Analyses

For testis histology, immunofluorescence in cancer cells, and proliferation assays, images were acquired using a 40X (testis histology and proliferation assays) or 60X (immunofluorescence in cancer cells and testis histology) objective on an inverted Nikon Ti2 AX-R confocal microscope with NIS-Elements software (Nikon). All images were analyzed using Fiji^120^. For testis histology, mean intensity was calculated for manually drawn ROIs of nuclei on grayscale, maximum-intensity projections and normalized to the background for each respective channel. For proliferation assays thirteen images per sample from three independent biological replicates were acquired, nuclei were quantified and averaged per replicate, then normalized to the scrambled control. For testis histology with c-KIT, images were acquired using a 20X objective on an Olympus Fluoview FV1000 confocal laser-scanning microscope (Olympus).

For migration assays, cell migration into the wound area was monitored using the Incucyte SX5 Live-Cell analysis system (Sartorius #4816) until ∼95-98% wound closure (∼8-20 h for MDA-MB-231 cells and ∼68-72 h for HCC-1954). Images were acquired every 2 h using a 10X objective and analyzed to calculate relative wound density, defined by the Incucyte software as the density of the wound region (%) relative to the density of the cell-occupied region, defined initially at the start of image acquisition. Three technical replicates were analyzed using the Incucyte Scratch Wound Analysis software module (Sartorious #9600-0012).

### Statistical Analyses and Graphs

Graphs for testis histology (**Supplementary Fig. 2C**), RT-qPCRs, migration and proliferation assays were generated using GraphPad Prism 10.2.3. for Mac OS X (GraphPad Software, Boston, FsMassachusetts USA, www.graphpad.com). All statistical analyses were performed using an unpaired, two-tailed Student’s t-test with Welch’s correction, unless otherwise as noted in the figure legends.

### Identification of ACTL8 Orthologs and Canonical Arps

True syntenic ACTL8 orthologs were collected by first identifying conserved flanking genetic neighbors in the human genome (version hg38)^121^: *ARHGEF10L*, *IGSF21,* and *RCC2*. These genes were identified in the sequenced and assembled genomes of other vertebrates listed in **Supplementary Data 1** using the University of California Santa Cruz (UCSC) genome browser (http://genome.ucsc.edu)^122^. *ACTL8*-like genes present between these flanking genes were collected and then verified by NCBI’s annotations (**Supplementary Data 1**) and phylogenetics (see Phylogenetics section). If genomes were not well annotated for an *ACTL8* gene, human ACTL8 or an ACTL8 protein sequence in a closely related species was used as a query in NCBI tBLASTn^49^ and then aligned to the genome in UCSC’s Genome Browser using the BLAT function^123^. Synteny was then verified by identification of the genetic neighbors *ARHGEF10L, IGSF21,* and *RCC2*. For platypus, tBLASTn initially identified an *ACTL8*-like gene, and synteny was confirmed using one of the latest genomes for platypus on the UCSC genome browser from 2020 (Pmale09 v4 2020, GCF_004115215.2). When conducting tBLASTn with invertebrate genomes, the top hits were other members of the Arp superfamily (**Supplementary Data 1**), consistent with our failure to detect *ACTL8* in the syntenic locus in the corresponding genome assembly on the UCSC Genome Browser.

Canonical Arps are generally well-annotated in mammalian genomes and were identified either by the UCSC genome browser or NCBI. Synteny was determined by verifying the presence of highly conserved flanking genes that were in common with the human genome. Syntenic neighbors and accession codes are listed in **Supplementary Data 1.**

### Sequence Alignments, Phylogenetics, and Structural Analyses

Species trees were generated using TimeTree5^50^. Protein sequences of ACTL8 orthologs with and without canonical Arps were aligned using MAFFT^124^ with the SnapGene software (www.snapgene.com). For maximum likelihood phylogenetic analyses, the alignment was uploaded to the IQ-TREE webserver^125^ and default parameters were used with selection of the LG substitution model for amino acid sequences^126^ and ultrafast bootstrap with 1,000 replicates for analyzing statistical support. The newick file generated by IQ-TREE was viewed using FigTree (version 1.1.4, https://tree.bio.ed.ac.uk/software/figtree/) and is provided in **Supplementary Data 3** with the corresponding protein alignment in **Supplementary Data 2**. AlphaFold 3.0^51^ was used to generate a predicted structure of human ACTL8 bound to ATP and Mg^2+^ as ligands. ChimeraX^127^ (version 1.7) was used to view the structure and map sequence conservation onto its surface. Conservation was viewed by importing MAFFT-generated sequence alignments and using the “Render by Attribute” tool.

Protein sequences for human KDM5 family members (**Supplementary Fig. 10**) were obtained from UniProt^128^. Alignments were performed using Clustal Omega^129^ and visualized in Jalview^130^. Pairwise sequence identities were calculated in Jalview or Snapgene.

### Yeast-Two-Hybrid Assay

Potential binding partners of ACTL8 were screened for via a yeast-two-hybrid assay, done by the company Hybrigenics Services (https://www.hybrigenics-services.com). The “prey” was a cDNA library from the human testis ([HTestis]), and the bait was full-length human ACTL8 (with original human codon usage) cloned into vector pB29. First, the LexA system was used with a C-terminal fusion of the LexA DNA-binding domain. Due to lack of toxicity of the ACTL8 construct, a selective medium without 3-Aminotriazol was used for the screens. Because only 3 positive clones were grown among 109 million tested interactions, the screen was switched to the more sensitive Gal4 system. For this system, the DNA-binding domain Gal4 was fused to the N-terminus of ACTL8 in the vector pB66. The pB66-Gal4-ACTL8 construct was not toxic or auto-activating, and the system yielded 380 positive clones out of 62 million tested interactions.

### Positive Selection Analyses

The PAML suite^59^ was used to test for statistical significance of positive selection. PAML’s codeml algorithm tests nucleotide alignments to determine whether a subset of residues evolved under positive selection, asking whether a dN/dS ratio >1 (positive selection) is more likely than neutral evolution. Here, dN represents the rate of nonsynonymous (amino-acid altering) changes, and dS is the neutral ’control’ rate of synonymous changes. We first collected ACTL8 orthologs from simian primates, because in our experience these species span a suitable evolutionary distance for PAML analysis. We used the human ACTL8 ORF (from Genbank NM_030812) as a query in blastn searches of NCBI’s nr database and downloaded the best match from each species. We also performed reciprocal blastn with each candidate ortholog against a database of all human ORFs to ensure that ACTL8 was the best hit, ie. to ensure we had obtained orthologs rather than paralogs. We generated an in-frame nucleotide alignment (**Supplementary Data 4**) using the MACSE v2 frame-aware alignment algorithm [ref https://pubmed.ncbi.nlm.nih.gov/30165589/]. We trimmed an N-terminal extension and unexpected internal 15 aa insertion near the splice junction from one predicted ortholog (Francois’s langur, Genbank XM_033228885) and estimated a phylogeny using the PHYML maximum likelihood algorithm, version 3.3.20220408 [ref https://pubmed.ncbi.nlm.nih.gov/20525638/], with the GTR substitution model, 4 rate categories, and estimating the proportion of invariant sites.

The alignment and tree were used as input files in PAML^59^, using codeml with the F3x4 codon frequency model (CodonFreq=2), initial omega 0.4, and retaining sites with alignment gaps (cleandata=0). We ran codeml with each of three evolutionary models (M7, M8a, and M8), and compared the resulting likelihoods to determine whether a model that allows for a subset of sites with dN/dS >1 (Model 8) is a better fit for the data than each of two different models that only allow sites under neutral and purifying selection (Model 7 or Model 8a). A chi-squared distribution was used to assess statistical significance of twice the difference in log-likelihoods of the models. Overall selective pressures were estimated using Model 0, which assumes a single dN/dS value for every site in the alignment; the same input files were used as before.

## Acknowledgments

We thank Angelique Whitehurst, Elisabeth Martinez, Jihan Osborne, and the Schroeder lab members for their comments on the manuscript. We are grateful to Jihan Osborne for the use of her Incucyte (funded by the Cancer Prevention and Research Institute of Texas RR210016) and to Bareun Kim for Incucyte training. We thank Angelique Whitehurst and Magdalena Delgado for providing cancer cell lines, constructs, and experimental suggestions. We also thank Elisabeth Martinez and Quin Zhang for discussions regarding KDM5B, and Jun Fang, Ram Madabushi, and Laura Banaszynski for technical advice on CUT&RUN and genomics analyses. We thank Kyle Orwig, Georgia Rae Atkins, and Kotaro Sasaki for discussions related to human sperm development and testis histology. We acknowledge UT Southwestern’s Genomics Core (Carlos Arana and Chaoying Liang) for conducting RNA sequencing and analysis and the McDermott Center Next Generation Sequencing Core (Vanessa Schmid) for CUT&RUN sequencing. Novogene and Plasmidsaurus provided additional sequencing services, and Hybrigenics Services conducted the yeast two-hybrid screen. This work was supported by the Cancer Prevention and Research Institute of Texas (RR210048 to C.M.S.), the National Institutes of Health (R35GM156807 and R00GM137038 to C.M.S.; R00GM138920 and R35GM166445 to S.Br.; R01HD110170 to C.B.G.; R01GM074108 funded J.M.Y. and I.M.N.), the UT Southwestern Endowed Scholars Program in Medical Science (C.M.S.), and a UT System Rising STARs award (C.M.S.).

## Author Contributions

N.C.S: methodology, data acquisition, data analysis, data curation, validation, writing, editing, and figure production. S.Ba.: data analysis, writing, editing and figure production for the CUT&RUN experiments. K.D.A.: data acquisition, data analysis, data curation for cytology and subcellular fractionation. S.P.: data acquisition, data analysis, data curation for testis histology. I.M.N.: initial data acquisition and data curation for phylogenetic analyses. B.C.: data acquisition and data curation for testis histology of c-KIT. C.B.G.: supervision of the testis histology performed by B.C. J.M.Y.: data acquisition and analysis for positive selection analyses. S.Br.: supervision of the CUT&RUN data analysis and editing. C.M.S.: conceptualization, project management, funding acquisition, methodology, data acquisition and analysis, phylogenetics, figure production, writing and editing. N.C.S. and C.M.S. wrote the manuscript with input from all authors.

---

**Supplementary Data 1:** Genomic analysis of ACTL8 orthologs and canonical Arps, related to Figure 1 and Supplementary Figure 1.

**Supplementary Data 2:** Alignment for phylogenetic analysis in Figure 1B.

**Supplementary Data 3:** Newick tree file for phylogenetic analysis in Figure 1B.

**Supplementary Data 4:** Alignment of primates sequences used for positive selection analyses in Supplementary Figure 1D.

**Supplementary Data 5:** List of ACTL8-Regulated Genes in MDA-MB 231 Cells RNA-seq knockdown and overexpression, related to Figure 3.

**Supplementary Table 1:** Antibodies used in this study.

**Supplementary Table 2:** RT-qPCR primers used in this study.

**Supplementary Table 3:** Plasmids generated in this study.

**Supplementary Movie 1:** Representative movie of MDA-MB-231 WT scrambled (negative control) in wound healing assay, related to Figure 4B.

**Supplementary Movie 2:** Representative movie of MDA-MB-231 ACTL8 knockdown (oligo 1) in wound healing assay, related to Figure 4B.

**Supplementary Movie 3:** Representative movie of MDA-MB-231 no Doxycycline treatment negative control in wound healing assay, related to Figure 7B.

**Supplementary Movie 4:** Representative movie of MDA-MB-231 PHD2-α2 induced overexpression in wound healing assay, related to Figure 7B.

**Supplementary Fig. 1:**
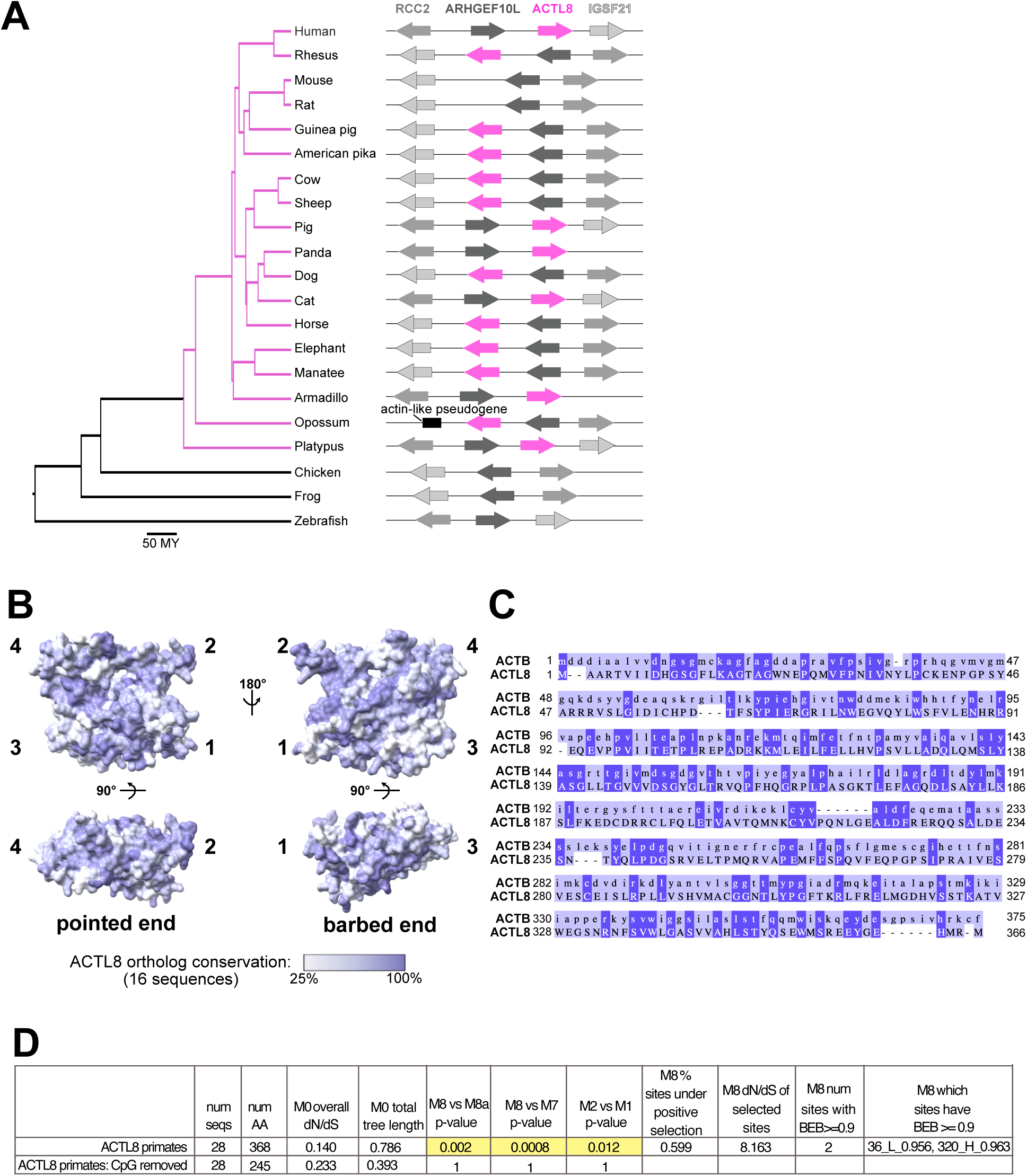
ACTL8 evolutionary analyses and evidence of positive selection in primates, related to. **Figure 1. A)** Schematic of the *ACTL8* locus among representative vertebrate species. Flanking genes *RCC2*, *ARHGEF10L*, and *IGSF21* were used to confirm synteny. Syntenic *ACTL8* orthologs were only found in mammals among the species surveyed. **B)** AlphaFold 3.0^51^ predicted structure of ACTL8 with conservation of all collected ACTL8 orthologs (16 sequences) mapped onto the surface of the structure. Subdomains and the pointed or barbed ends as found in actin are noted, and conservation is shown as a gradient with 100% conservation in blue and lowest conservation (25%) in white. **C)** Protein sequence alignment of human ACTL8 and actin (ACTB). Amino acid identities are indicated by color (100% identical residues are shaded dark blue; unconserved residues are shaded light blue). **D)** Tables showing the results of positive selection analyses with PAML^59^, using primate sequences with and without a CpG mask (see methods). Tree length indicates the number of nucleotide substitutions for each codon. Statistically significant values are highlighted, indicating ACTL8 evolves under positive selection. However, when CpG sites are not considered, signal for positive selection is not statistically significant. The M8 model indicates two positively selected sites with M8 Bayes Empirical Bayes (BEB) poster probability >90%.

**Supplementary Fig. 2:**
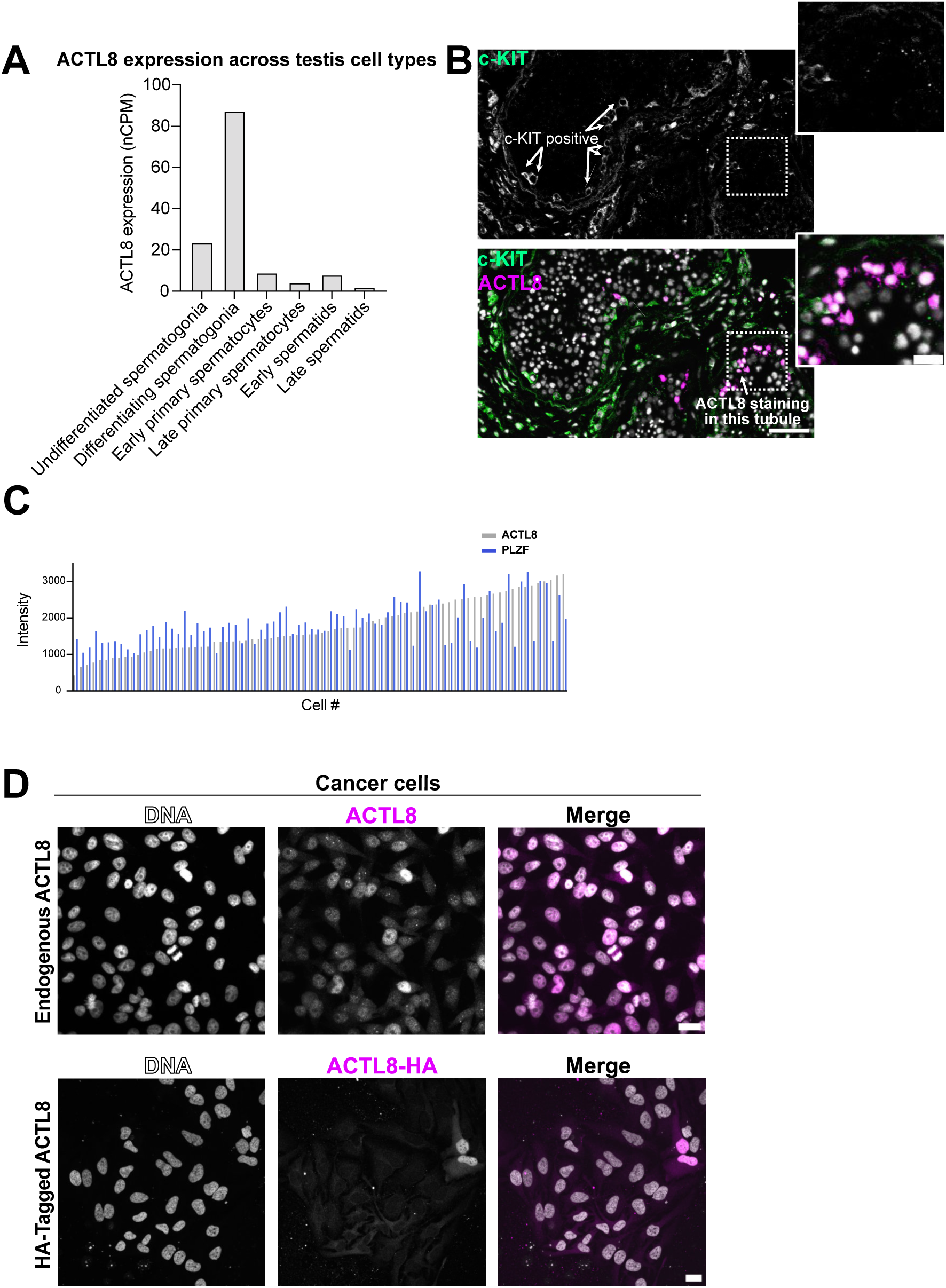
Quantification of ACTL8 and early spermatogenesis markers, related to. **Figure 2. A)** ACTL8 expression levels across human testis germ cell types from publicly available single-cell RNA-seq data^35,55^. **B)** Co-immunostaining of ACTL8 and c-KIT (differentiated spermatogonia). Left: whole seminiferous tubules. Arrows indicate c-KIT-positive and ACTL8-positive cells. The tubule on the left displays multiple c-KIT-positive cells but no detectable ACTL8-positive cells. The tubule on the right displays fewer c-KIT-positive cells and multiple ACTL8-positive cells. No strong colocalization between ACTL8 and c-KIT was observed in the tubule on the right. Scale bar: 100 µm. Right: higher magnification image of ACTL8-positive and c-KIT-positive cells from the rightmost tubule, scale bar: 30 µm. **C)** Quantification of ACTL8 and PLZF intensities per cell. Each bar represents a single cell. Data were obtained from four seminiferous tubules. **D)** Lower magnification of ACTL8 immunofluorescence in cancer cells. Top: endogenous ACTL8 in MDA-MB-231 cells using the anti-ACTL8 antibody, as in Fig. 2D. Bottom: HeLa cells expressing exogenously expressed HA-tagged ACTL8, stained with an anti-HA antibody. Scale bar: 20 µm.

**Supplementary Fig. 3:**
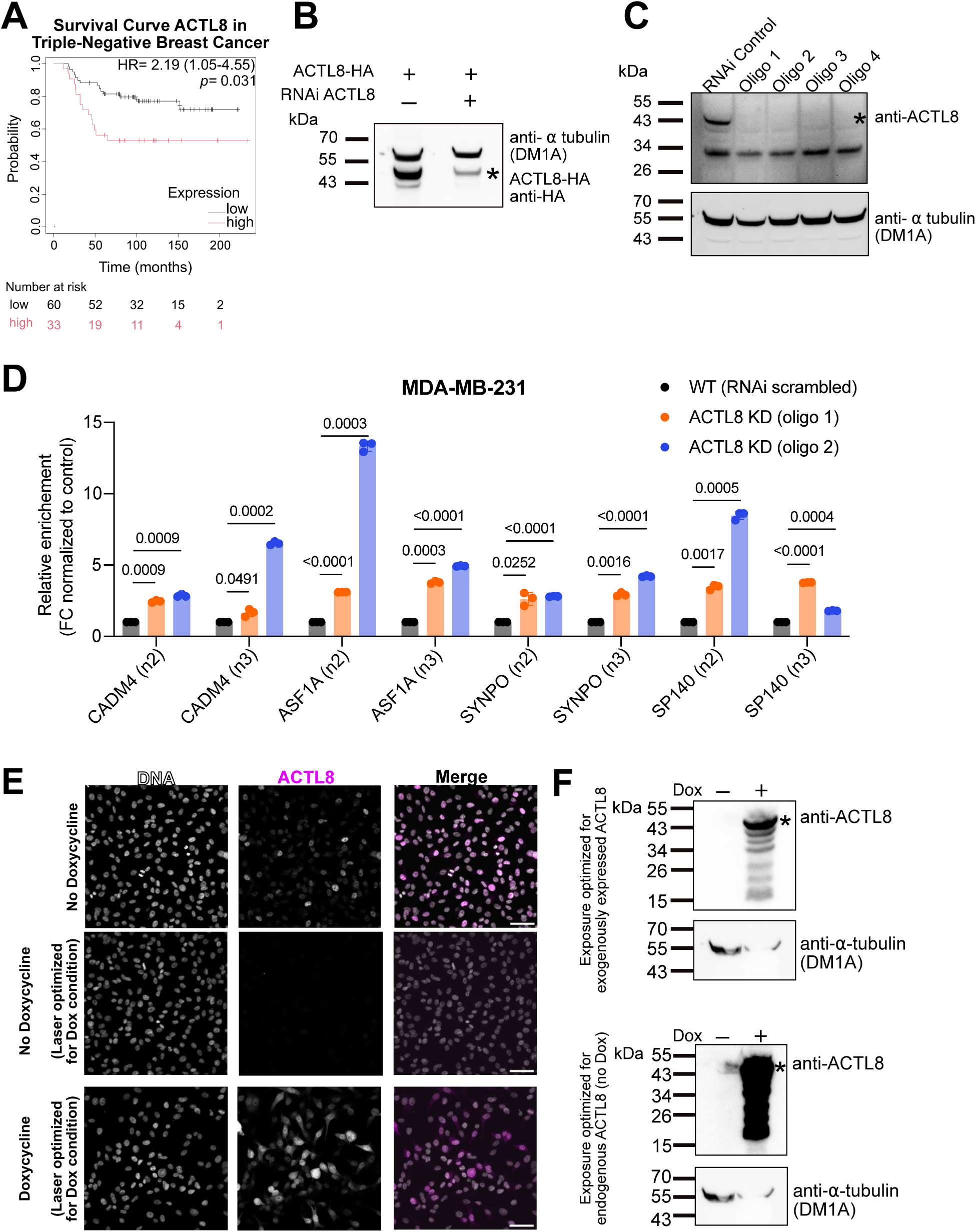
Validation of *ACTL8* knockdown, transcriptional effects, and doxycycline-inducible overexpression in MDA-MB-231 cells, related to. **Figure 3. A)** Survival curve of ACTL8 expression in breast cancer^79^ were generated using the Kaplan–Meier Plotter^134^ database. Graph acquired in January 2025. **B)** Immunoblot of HEK-293T cells expressing exogenously expressed ACTL8-HA, probed with an anti-HA antibody. Cells were treated with control siRNA or *ACTL8*-targeting siRNA. ACTL8-HA (indicated with an asterisk) is reduced upon *ACTL8* knockdown, an additional control confirming siRNA knockdown as previously assessed with an anti-ACTL8 antibody. Tubulin was used as a loading control. Anti-HA and anti-tubulin were probed sequentially. **C)** Immunoblot showing efficient knockdown of *ACTL8* in MDA-MB-231 cells using scrambled siRNA (RNAi control) or individual siRNAs (four independent siRNAs comprising the siRNA pool targeting *ACTL8* mRNA). The ACTL8 band is indicated by an asterisk. Tubulin was used as a loading control. **D)** RT–qPCR validation of selected genes from the RNA-seq dataset (see Figure 3B), showing two additional independent biological replicates that show similar changes in gene expression as seen in Figure 3C. Each dot represents a technical replicate (n = 3). Expression levels were normalized to 18S rRNA across samples, and the relative expression of oligo 1- and oligo 2-treated samples was normalized to the WT (scrambled siRNA) condition. Statistical analysis was performed using an unpaired t-test with Welch’s correction. *ACTL8* knockdown was performed using two independent siRNAs as in Fig. 3C. **E)** Exogenously expressed *ACTL8* was assessed by immunofluorescence in MDA-MB-231 cells infected with adenovirus and cultured in the presence or absence of doxycycline, which induces expression of *ACTL8*. Images were acquired either with laser settings optimized for each condition or with image acquisition settings optimized for the overexpression of *ACTL8* and kept constant across all samples. Relative levels show endogenous ACTL8 is present, but overexpression of *ACTL8* leads to much higher levels of protein. Scale bar: 50 µm. **F)** Immunoblot of MDA-MB-231 cells infected with adenovirus and incubated with or without doxycycline to drive *ACTL8* overexpression. Membranes were probed with anti-ACTL8 antibody (indicated by an asterisk). Top: Exposure conditions were optimized to visualize exogenously expressed ACTL8, resulting in no detectable endogenous ACTL8 signal in the condition without doxycycline. Bottom: Exposure conditions were optimized to visualize endogenous ACTL8 in absence of doxycycline, leading to saturation of signal in the condition with doxycycline. Because both images were acquired from the same membrane, the same tubulin loading control is shown in the top and bottom panels.

**Supplementary Fig. 4:**
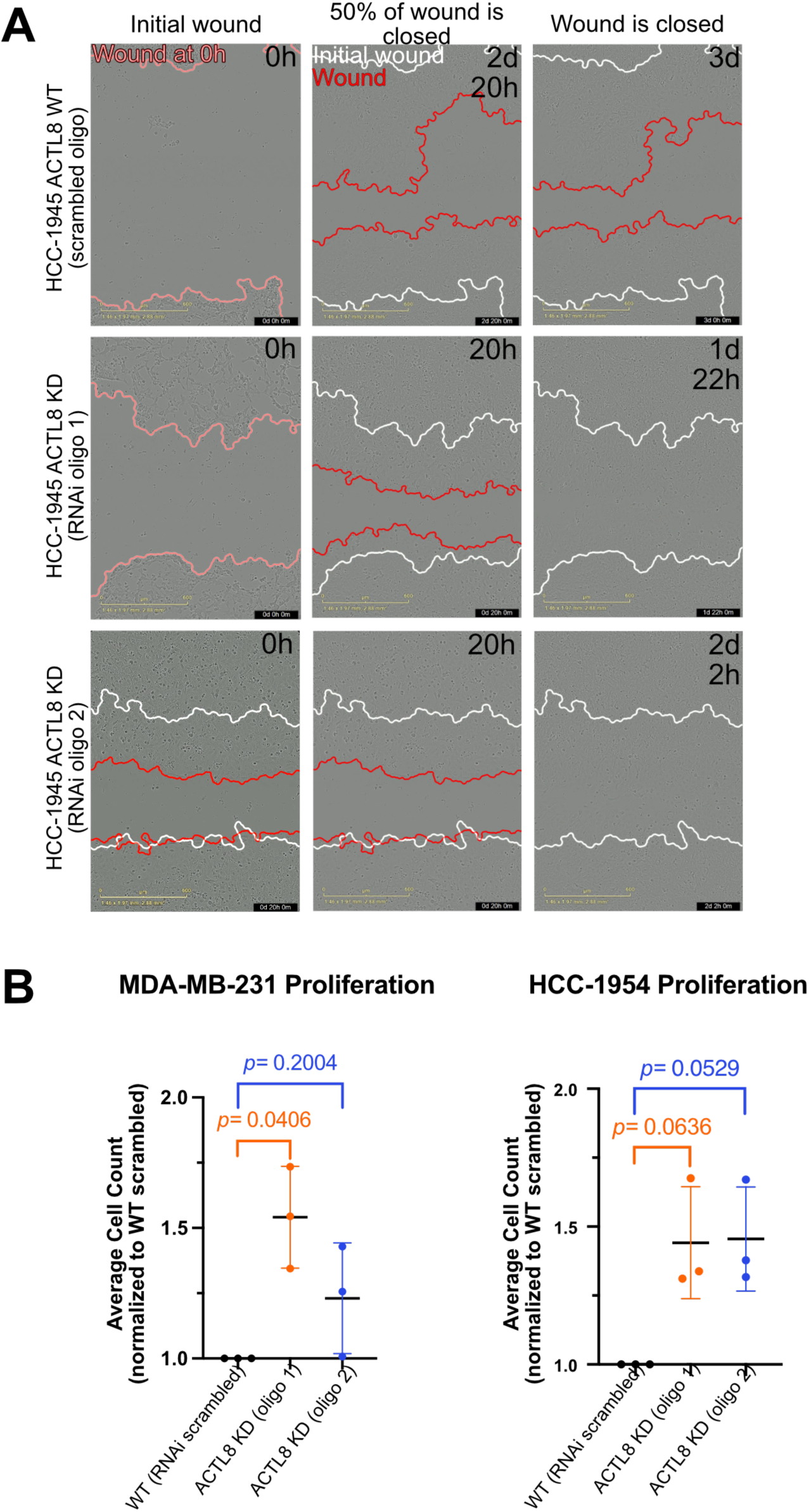
Representative images of HCC-1954 cells in wound healing assay and quantification of cell proliferation changes upon *ACTL8* knockdown, related to. **Figure 4. A)** Representative images of three time points from live-cell wound-healing assays in HCC-1954 cells transfected with scrambled siRNA (WT) or *ACTL8*-targeting siRNAs (“oligo 1” and “oligo 2”). Images show the initial wound (0 h), ∼50% wound closure, and wound closure. The time required to reach each stage is indicated in the top right of each image. **B)** Quantification of cell proliferation, measured as number of nuclei, in MDA-MB-231 and HCC-1954 cells treated with siRNA scrambled (WT, in black) or two independent *ACTL8*-targeting siRNAs. Following knockdown, cells were fixed and stained with a DNA probe. Nuclei were imaged and quantified using Fiji^120^. Datapoints represent three independent biological replicates, each performed with 13 images per sample and averaged per replicate, then normalized to the scrambled control. Statistical significance was evaluated using an unpaired, two-tailed Student’s t-test with Welch’s correction.

**Supplementary Fig. 5:**
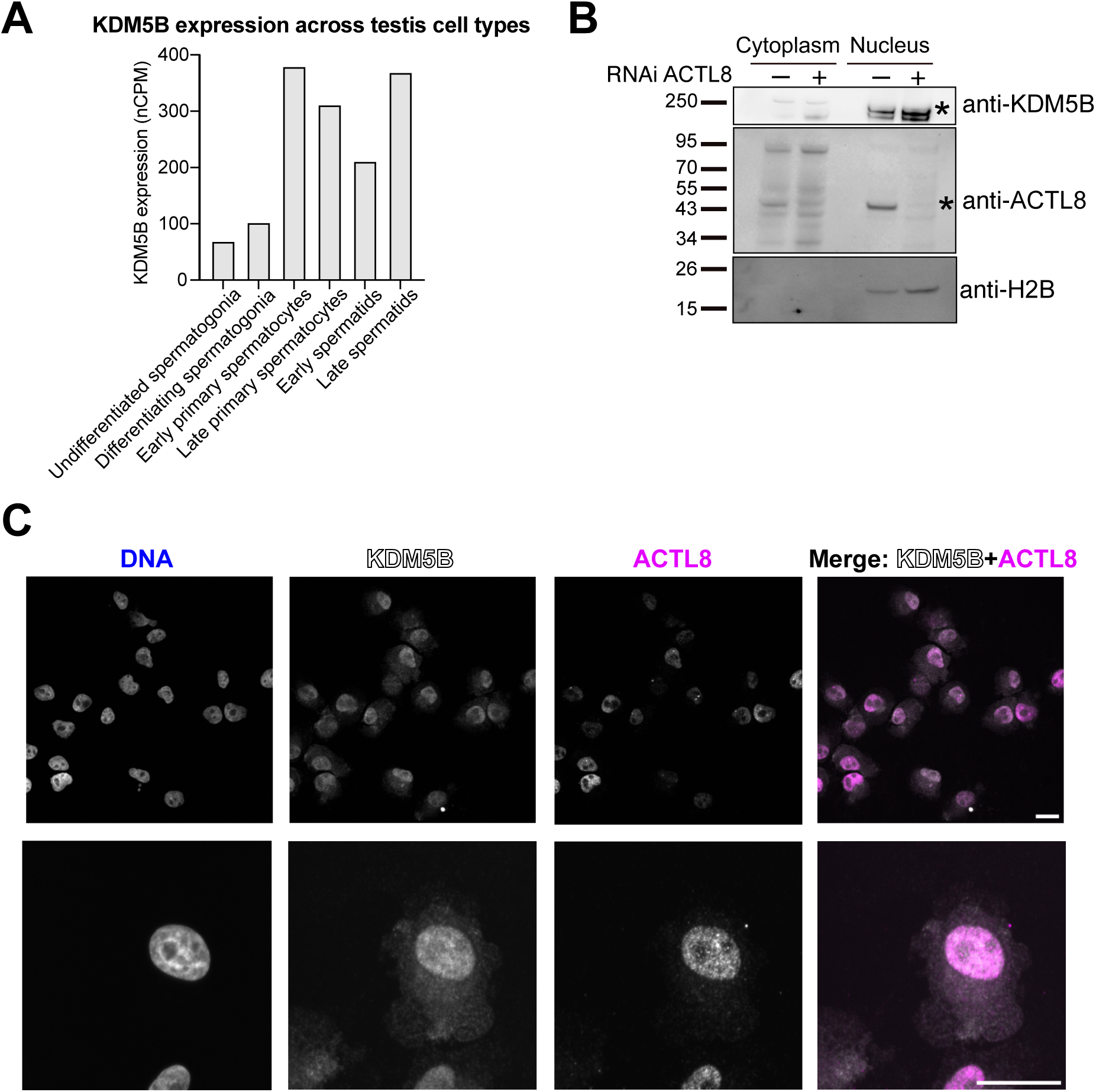
Expression levels and localization of KDM5B, related to. **Figure 5. A)** KDM5B expression levels across human testis germ cell types from publicly available single - cell RNA-seq data^35,55^. **B)** Immunoblot of subcellular fractionation in MDA-MB-231 cells under control conditions (siRNA scrambled) or upon *ACTL8* knockdown (siRNA *ACTL8* pool), probed with anti-KDM5B and anti-ACTL8 antibodies. Histone H2B was used as a chromatin-bound marker. **C)** Co-immunostaining of endogenous KDM5B and ACTL8, showing co-localization in the nucleus. Top: lower-magnification view. Bottom: higher-magnification view. Scale bar: 20 µm.

**Supplementary Fig. 6:**
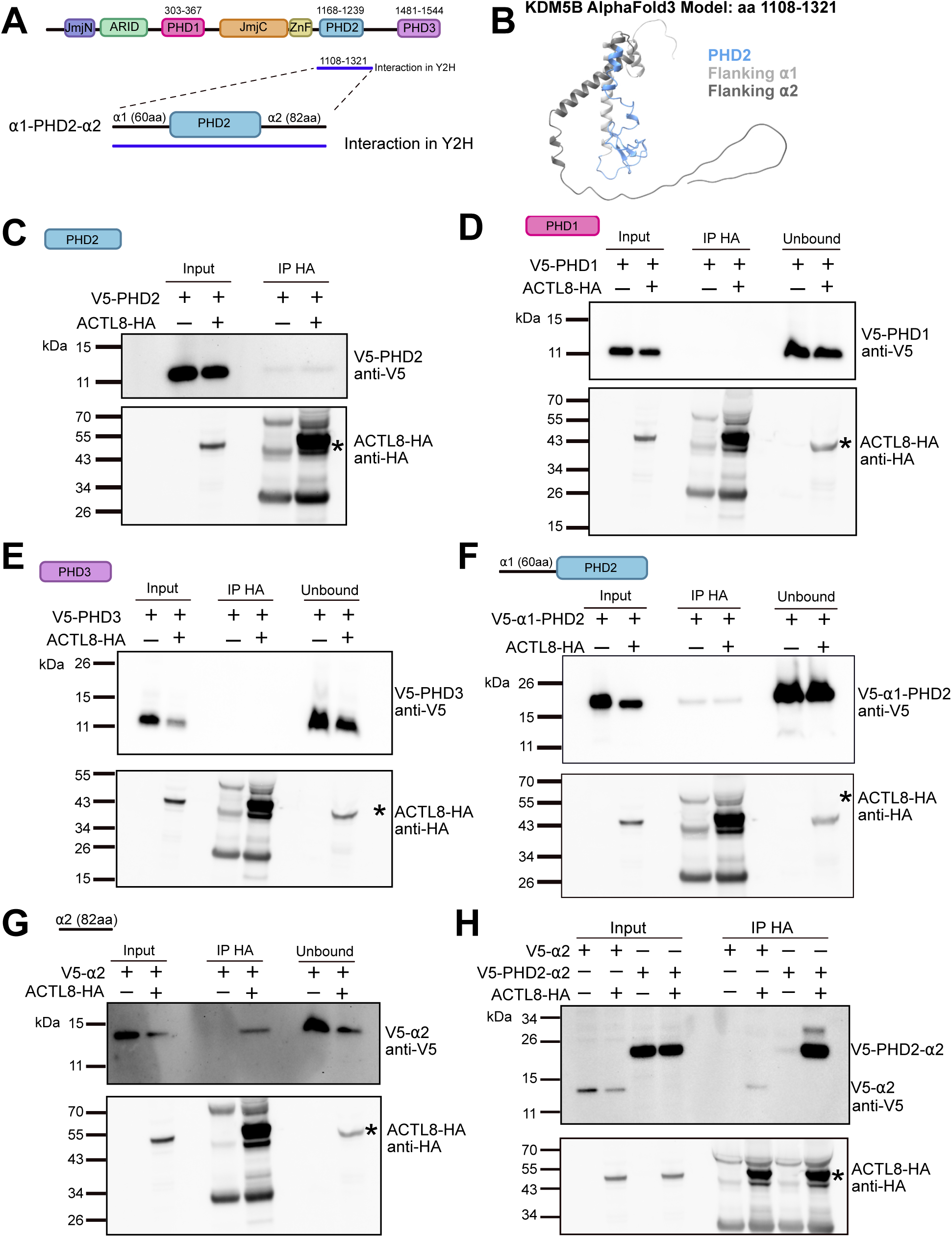
Mapping the KDM5B region interacting with ACTL8, related to. **Figure 5. A)** Schematic of KDM5B domain structure (amino acid positions were assigned based on Klein et al., 2014^86^), highlighting the ACTL8-interacting region identified in the yeast two-hybrid (Y2H) screen. A zoom-in of this region shows two *a*-helices flanking the PHD2 domain that form the “*a*1-PHD2” or “PHD2-*a*2” fragments tested in panel F and Figure 5D. Amino acid positions corresponding to the PHD domains and the Y2H are indicated. **B)** AlphaFold 3.0^51^ predicted structure of KDM5B. Only displaying the Y2H fragment with the PHD2 domain and flanking *a*-helices distinguished by color. **C-G)** Immunoblot of a co-immunoprecipitation (co-IP) with HEK-293T cells expressing ACTL8-HA and the indicated V5-tagged KDM5B fragment, either PHD2, PHD1, PHD3, *a*1-PHD2 or *a*2 alone. In all co-IPs, ACTL8-HA was immunoprecipitated using anti-HA magnetic beads and samples were probed for HA and V5 tags. Asterisk indicates the band corresponding to ACTL8, only the α2 alones interact with ACTL8, albeit weakly compared to the PHD2-α2 fragment (see Fig. 5D). **H)** Immunoblot of a co-IP as in C-G *a*2 alone versus PHD2-*a*2 were compared. PHD2-*a*2 expresses better and appears to bind more robustly than *a*2 alone.

**Supplementary Fig. 7:**
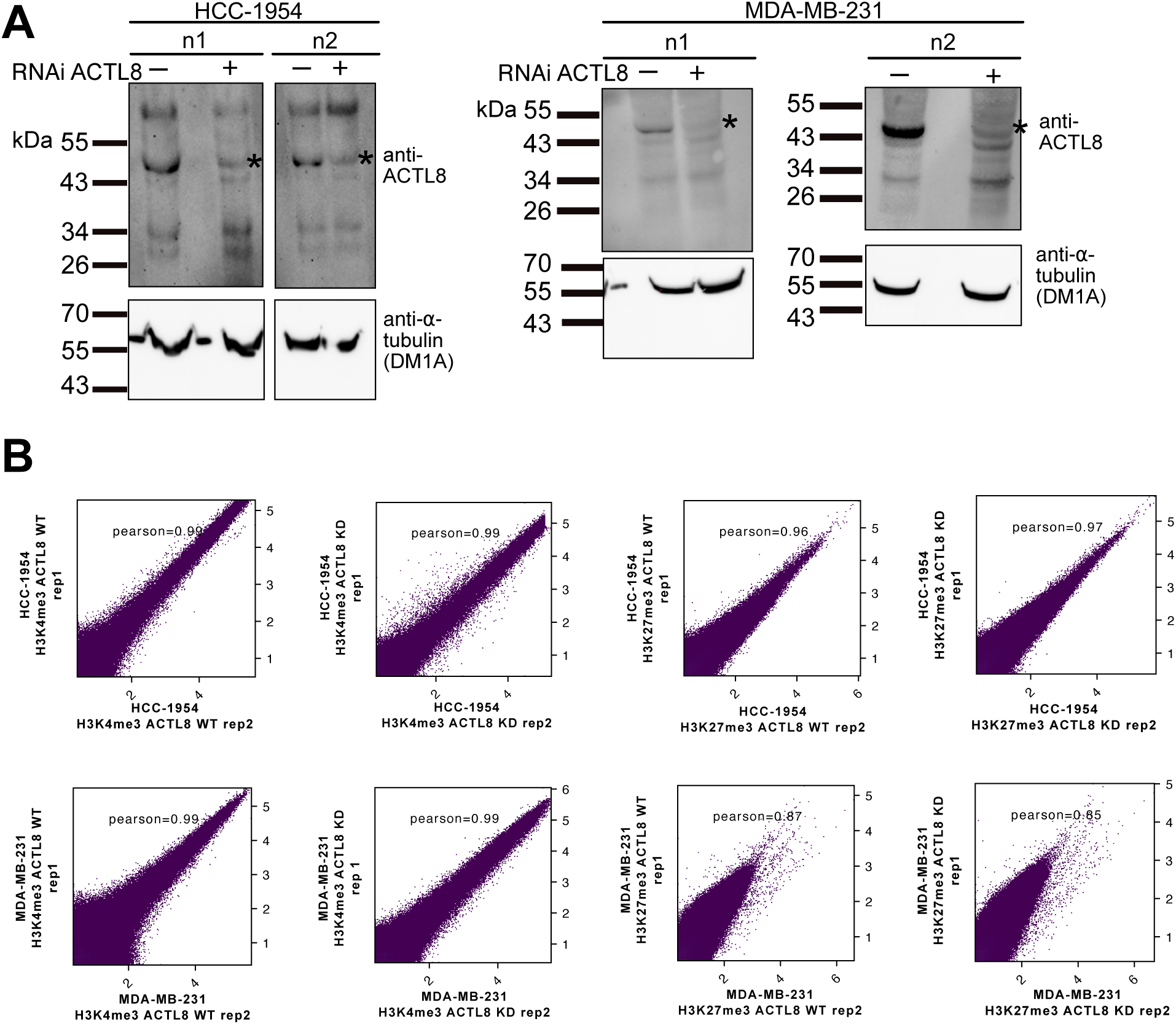
Validation of *ACTL8* knockdown and reproducibility of CUT&RUN datasets, related to. **Figure 6. A)** Immunoblots showing efficient knockdown (KD) of *ACTL8* in HCC-1954 and MDA-MB-231 cells using an *ACTL8*-targeting siRNA pool in the two biological replicates (n1, n2) used for CUT&RUN assays (see Fig. 6). The ACTL8 band is indicated by an asterisk, and tubulin was used as a loading control. **B)** Scatter plots comparing genome-wide signal between biological replicates for H3K4me3 and H3K27me3 in *ACTL8* WT and KD conditions across HCC-1954 and MDA-MB-231 cell lines. Each point represents a genomic bin, plotted as log-transformed signal intensity in replicate 1 versus replicate 2. Color density reflects point concentration. Pearson correlation coefficients (r) are indicated for each comparison, demonstrating high reproducibility between replicates.

**Supplementary Fig. 8:**
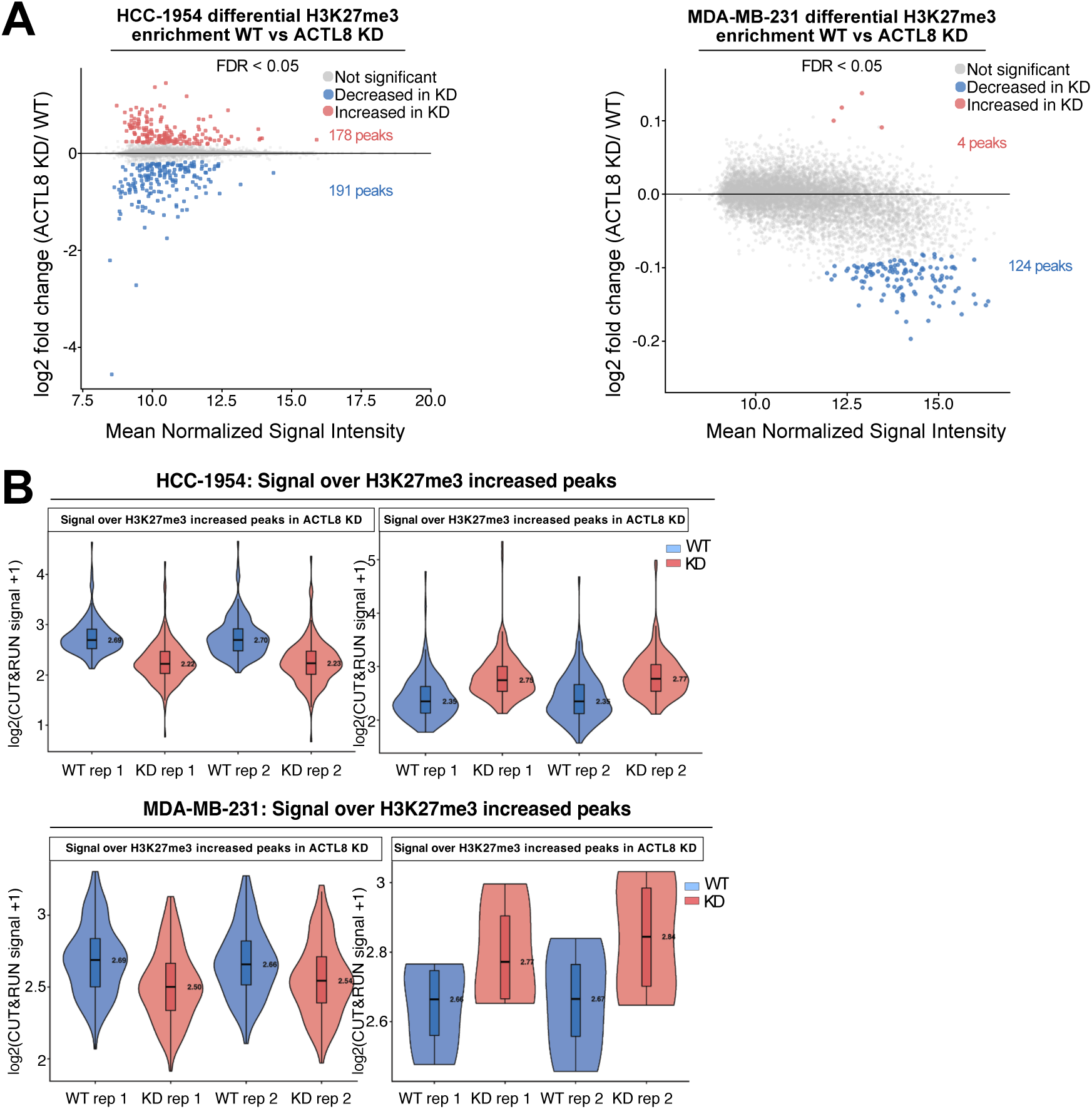
H3K27me3 shows limited changes upon *ACTL8* knockdown, related to. **Figure 6. A)** MA plots showing differential H3K27me3 enrichment between *ACTL8* WT (scrambled siRNA) and *ACTL8* knockdown (KD) in HCC-1954 (left) and MDA-MB-231 (right) cells. Peaks were identified by DiffBind^117^, and differential enrichment was determined using a false discovery rate (FDR) < 0.05. Each point represents an H3K27me3 peak plotted by mean normalized signal versus log₂ fold change (KD/WT). Red and blue points denote peaks with increased or decreased H3K27me3 in *ACTL8* knockdown (KD), respectively, while grey points indicate peaks with non-significant difference. **B)** Violin plots showing H3K27me3 CUT&RUN signal over *ACTL8* KD-increased and *ACTL8* KD-decreased regions were plotted for HCC-1954 (top) and MDA-MB-231 (bottom) cells. KD-decreased regions exhibit higher H3K27me3 signal in WT samples than in *ACTL8* KD samples, whereas *ACTL8* KD-increased regions show higher signal in KD samples, validating the differential peak classification. Signal is shown as log₂ (CUT&RUN signal + 1) for two biological replicates, with boxes indicating the interquartile range and median values annotated within each violin.

**Supplementary Fig. 9:**
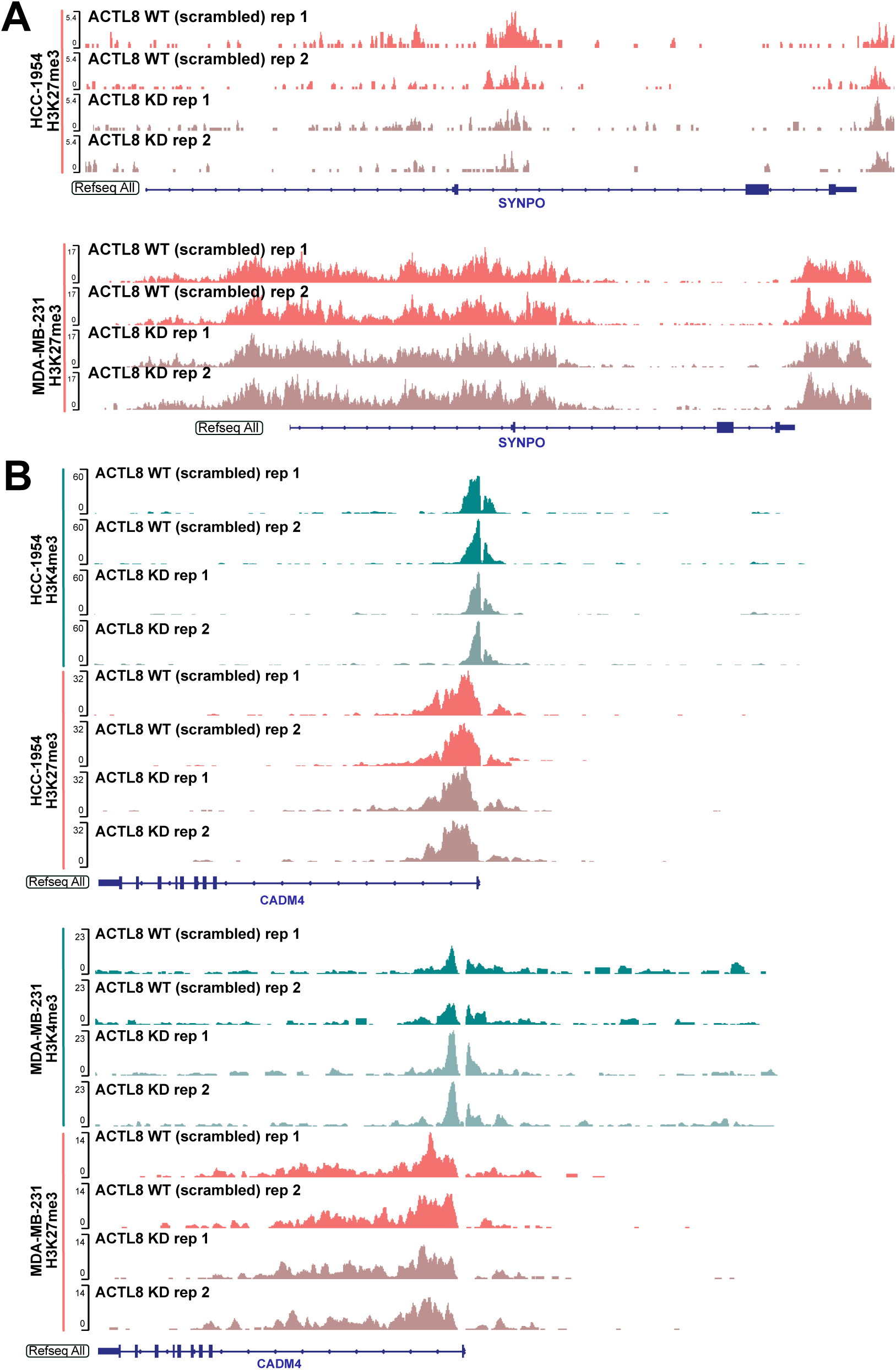
Additional IGV tracks from CUT&RUN experiments, related to. **Figure 6. A)** IGV tracks of *SYNPO* locus showing H3K27me3 occupancy. H3K27me3 is reduced in HCC-1954 cells with a slight reduction in MDA-MB-231 cells. **B)** IGV tracks of a representative locus that exhibits different H3K4me3 signals between the cell lines. *CADM4* shows a modest decrease in H3K4me3 and no obvious change in H3K27me3 in HCC-1954 cells, whereas *CADM4* shows an increase in H3K4me3 and no obvious change in H3K27me3 in MDA-MB-231 cells.

**Supplementary Fig. 10:**
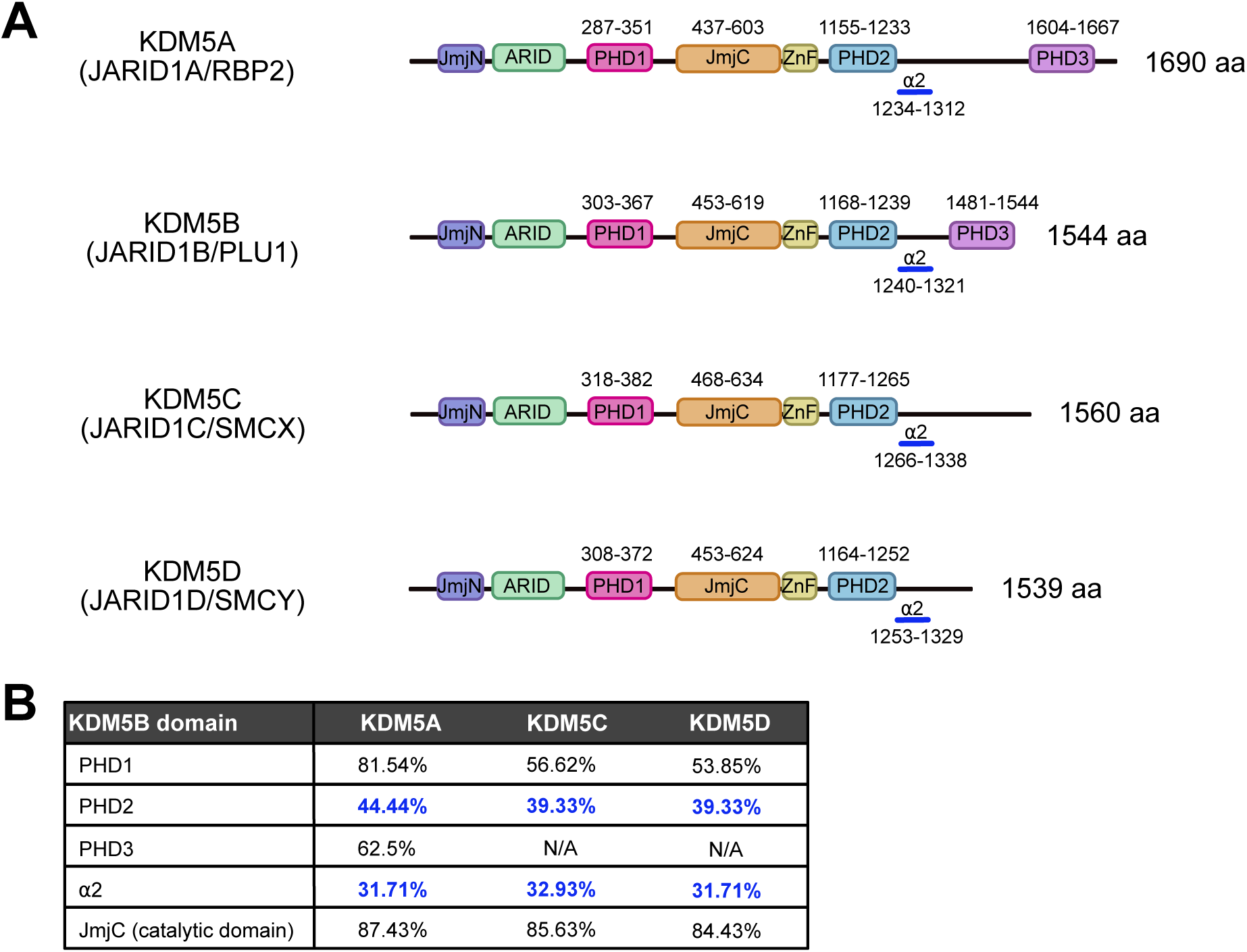
Sequence identity of KDM5B PHD fingers relative to other human KDM5 proteins. **A)** Schematic of KDM5B domain structure (amino acid positions were assigned based on Klein *et al.,* 2014^86^) and other human KDM5 proteins. The location of the catalytic domain and PHD fingers in KDM5A, KDM5C and KDM5D was inferred based on sequence alignment to KDM5B. KDM5C and KDM5D lack a PHD3 domain^83^. **B)** Pairwise sequence identify of the catalytic domain and PHD fingers across human KDM5 proteins. N/A indicates the absence of PHD3 fingers. Pairwise sequence identity was calculated from Clustal Omega^129^ alignments visualized in Jalview^130^.

